# Cytokine signaling regulates multiple stages of gammaherpesvirus infection in myeloid cells

**DOI:** 10.64898/2026.09.03.749271

**Authors:** Rachael E. Kostelecky, Gabrielle Vragel, Pedro Gamez, Eva M. Medina, Darby G. Oldenburg, Andrew Goodspeed, Christoph Weigel, Robert A. Baiocchi, Eric T. Clambey, Linda F. van Dyk

**Affiliations:** Department of Immunology and Microbiology, Anschutz Medical Campus, School of Medicine, Aurora, CO 80045 USA; Department of Anesthesiology, Anschutz Medical Campus, School of Medicine, Aurora, CO 80045 USA; Department of Biochemistry and Molecular Genetics, University of Colorado Denver Anschutz Medical Campus, School of Medicine, Aurora, CO 80045 USA; Gundersen Medical Foundation, Emplify Health System, La Crosse, WI 54601, USA; Department of Biomedical Informatics, | Anschutz Medical Campus, School of Medicine, Aurora, CO, 80045, USA; University of Colorado Cancer Center, University of Colorado Denver | Anschutz Medical Campus, School of Medicine, Aurora, CO, 80045, USA; Division of Hematology, Department of Internal Medicine, The Ohio State University, Columbus, OH 43210, USA

**Author notes:** Lead contact Linda F. van Dyk.

## Abstract

The gammaherpesviruses, including Epstein-Barr Virus and Kaposi’s Sarcoma-associated Herpesvirus, establish lifelong infections by maintaining a balance between lytic and latent infection states. How this balance is regulated in myeloid cells, an important but understudied cell type that can achieve both infection states, remains unclear. Using the murine gammaherpesvirus 68 (MHV68) model, we show that IFNγ and IL-4 reciprocally regulate lytic infection in macrophages in a time- and JAK/STAT signaling dependent-manner, with basal JAK/STAT signaling further restricting lytic infection. Using a combination of MHV68 reporter viruses, cellular and molecular techniques, we demonstrate that IFNγ blocks lytic replication through at least two distinct mechanisms, restricting delivery to or stability of the viral genome in the nucleus and potently restricting lytic cycle progression downstream of the immediate-early viral transactivator, RTA. These findings define specific points during lytic infection where IFNγ to enforce restricted, latent-like infection in myeloid cells.

**Graphical Abstract:** 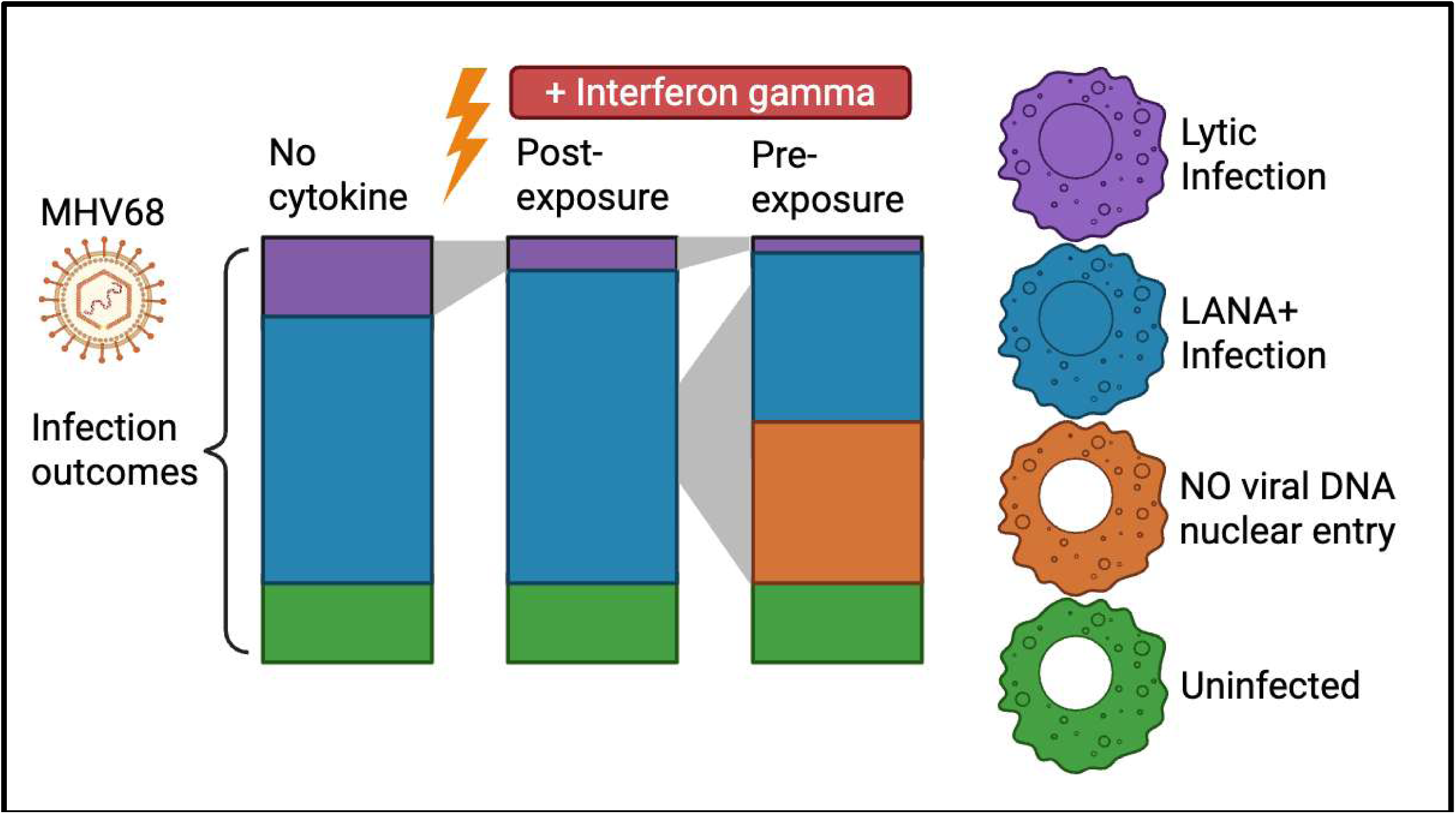

**Highlights:**

- IL-4 and IFNγ reciprocally regulate MHV68 macrophage infection in a time-dependent manner
- Lytic replication is induced by the JAK1/JAK2 inhibitor Ruxolitinib
- IFNγ regulates MHV68 infection at two distinct stages
- IFNγ pre-treatment limits viral nuclear entry

## INTRODUCTION

Gammaherpesviruses (γHVs) are DNA tumor viruses that include the human pathogens Epstein-Barr virus (EBV) and Kaposi’s sarcoma-associated herpesvirus (KSHV) and are associated with malignancies and inflammatory diseases^1,2,3^. γHV infection is controlled by the immune system, with decreased immune function resulting in increased risk of virus-associated disease^4^. The infection cycle of all γHVs is biphasic, characterized by an acute lytic phase, where viral genes are expressed in a coordinated cascade with viral DNA replication and production of new infectious virus particles, and a chronic latent phase where viral genomes persist in host cells with limited viral gene expression^4–6^. Reactivation from latency reinitiates productive lytic replication. The balance between lytic and latent states is shaped by host immune signaling, including cytokines.

The murine γHV model, murine gammaherpesvirus 68 (MHV68 or γHV68; ICTV nomenclature, *Murid herpesvirus 4*, MuHV-4), has genetic and biologic conservation with EBV and KSHV^2,4^, with lytic and latent infection states well characterized in epithelial cells and B lymphocytes, respectively^7^. MHV68 infection has also been demonstrated in peritoneal, alveolar, and splenic macrophages^2,8–11^. However, despite being an early target of infection in vivo^8,10^, myeloid cell infection remains poorly understood. MHV68 infection in myeloid cells links epithelial and B cell infection, where B cell infection significantly increases in the presence of macrophages^10^. Additionally, MHV68 infection of macrophages results in divergent infection states, where the majority of infected cells exhibit a latent-like, restricted infection characterized by limited viral gene expression, and only a small fraction of cells exhibit lytic infection^12^.

Macrophages are responsive to local immune environments and comprise diverse functional states that can be shaped by cytokines and other extracellular signals^13^, to alter infection outcomes^14,15^. Among these, interleukin-4 (IL-4) promotes lytic MHV68 replication through STAT6-dependent signaling^16,17^. In contrast, type II interferon (IFN), interferon gamma (IFNγ) promotes the establishment of latency, limits reactivation^18–20^, and prevents acute and chronic disease^21–24^ in MHV68-infected hosts. While the mechanisms by which IL-4 and IFNγ regulate infection remain under investigation, transcriptional regulation of the immediate early viral transactivator, RTA (encoded by *Orf50*), the driver of lytic replication^25^, appears to be a common point of regulation, with IL-4 inducing, and IFNγ repressing, RTA transcription^16,26^. The RTA promoter is further controlled by dynamic changes in epigenetic regulation, including histone acetylation and DNA methylation^27–31^. Enforced RTA expression enhances lytic replication and prevents latent infection^32^. How IL-4 and IFNγ regulate the balance between lytic and latent infection at the population- and single cell-level remains incompletely understood.

Here, we investigated how MHV68 infection of macrophages is regulated by cytokine signaling, integrating a suite of reporter viruses with cellular and molecular analysis of infection. We report that cytokine signaling reciprocally regulates replication in MHV68-infected macrophages, with IL-4 promoting and IFNγ restricting lytic replication with distinct temporal effects, and JAK/STAT signaling profoundly limiting lytic replication. We further identify that IFNγ is capable of restricting virus infection in at least two distinct phases of infection, with distinct consequences following IFNγ exposure before or after infection. Together, these findings identify multiple levels of IFNγ-dependent control of γHV infection in macrophages and emphasize the key role that JAK/STAT signaling has in limiting lytic replication in macrophages.

## RESULTS

### MHV68 infection of myeloid cells induces an interferon-inducible transcriptional signature in vitro and in vivo

We recently reported that MHV68 can efficiently infect macrophages, resulting in a high frequency of cells characterized by a restricted state of virus transcription, with few cells progressing to lytic replication^12^. How cytokine-dependent signaling influences these outcomes is unknown. To understand the transcriptional landscape of MHV68 infection in macrophages, we measured key virus (*Rta*) and host genes known to be induced by interferon (*Gbp5)* or IL-4 signaling (*Arg1*), in MHV68-infected J774 macrophages or in primary peritoneal macrophages isolated from infected C57BL/6J mice. In J774 cells, MHV68 infection resulted in induction of *Rta* (**Fig 1A**), consistent with initiation of lytic replication in a subset cells, with a modest induction of the interferon-inducible gene *Gbp5* (**Fig 1B**) and no discernible induction of the IL-4-inducible gene *Arg1* (**Fig 1C**). In parallel, we analyzed host gene expression in primary peritoneal macrophages isolated from immunocompetent C57BL/6J mice infected with the wild-type (WT) MHV68.LANA::βlac virus, a recombinant virus that allows direct identification and purification of virally-infected cells expressing the LANAβlac fusion protein, expressed during both lytic and latent infection^33^. At 16 hours post-infection (hpi), peritoneal macrophages positive for the macrophage marker F4/80 were FACS-purified into LANAβlac+ (LANA+) and LANAβlac-(LANA-) subsets and subjected to scRNA-seq (with quality control metrics in **Fig S1**). As expected, both populations expressed high transcripts for *Adgre1*, the gene which encodes the F4/80 protein (**Fig 1D**). When we examined IFN and IL-4 inducible transcripts, we found that LANA+ cells had increased expression of multiple IFN-inducible transcripts (*Isg15, Ifi204, Stat1,* **Fig 1E-G**) relative to LANA-cells isolated from the same animals, with lower expression of *Gbp5* and *Arg1* in both cell subsets (**Fig 1H-I**). These data indicate that de novo MHV68 macrophage infection is associated with induction of an early transcriptional signature consistent with interferon signaling.

**Figure 1.**
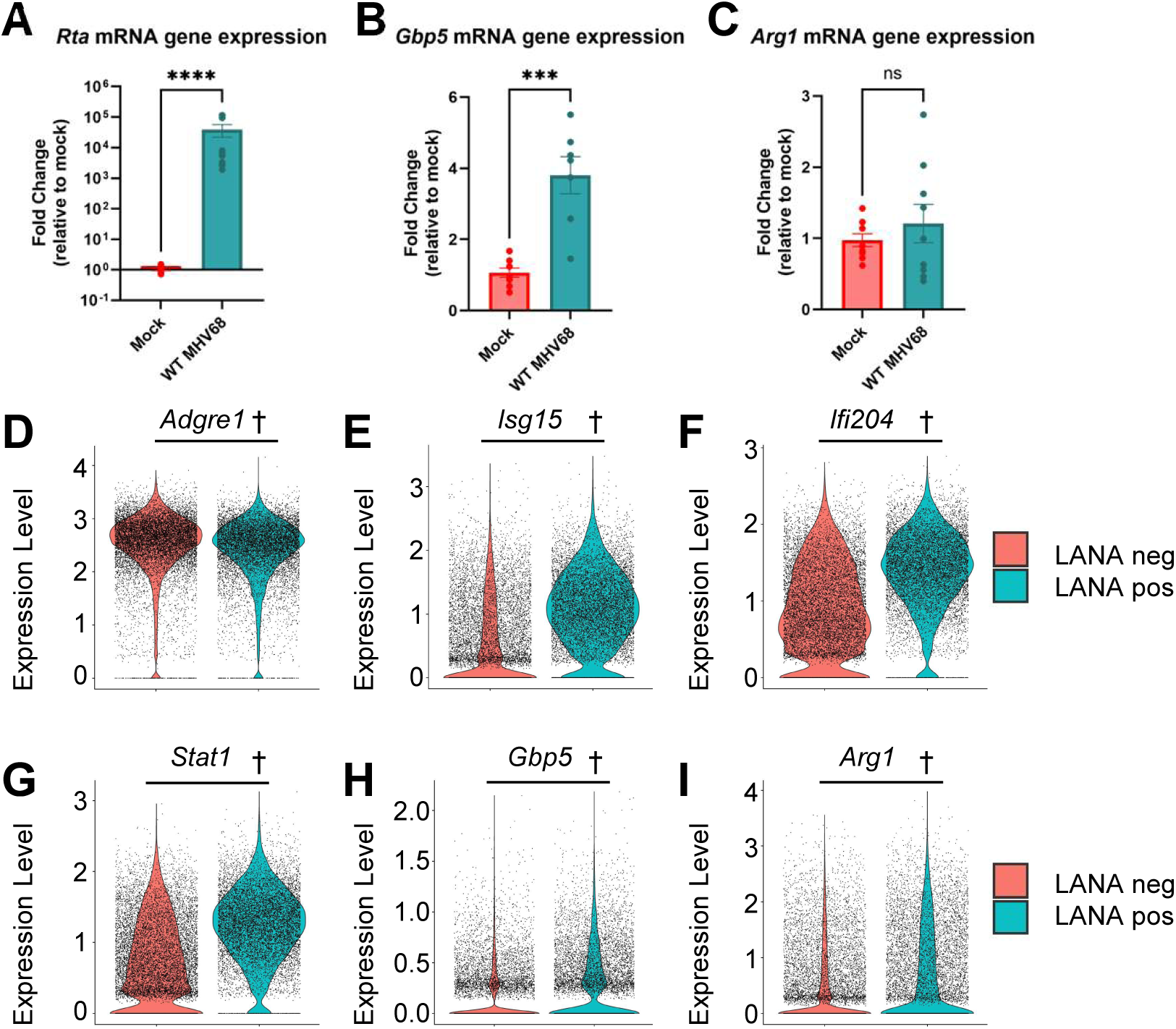
MHV68 infection of myeloid cells induces an interferon-inducible transcriptional signature in vitro and in vivo. Analysis of RNA expression in MHV68-infected **(A-C)** J774 macrophages or **(D-I)** primary peritoneal macrophages. **(A-C)** J774 cells were mock- or MHV68-infected at MOI=1 PFU/cell, harvested at 48 hpi, with RNA expression quantified by RT-qPCR, measuring **(A)** MHV68 *Rta*, encoded by *ORF50,* and cellular **(B)** *Gbp5*, and **(C)** *Arg1.* **(D-I)** C57BL/6J mice were infected with 1x10^6^ PFU of MHV68.LANA::βlac virus, with viable, F4/80+, LANA.βlac- and LANA.βlac+ peritoneal macrophages sort-purified at 16 hpi and subjected to scRNA-seq analysis. Violin plots, with data from individual cells indicated by circles, compare uninfected, LANA.βlac- and infected, LANA.βlac+ peritoneal macrophages, quantifying cellular RNAs for (D) *Adgre1*, which encodes for the F4/80 protein, **(E)** *Isg15*, **(F)** *Ifi204*, **(G)** *Stat1*, **(H)** *Gbp5*, **(I)** *Arg1*. Data **(A-C)** are from three independent experiments, with each experiment done in three parallel culture wells plated from a single culture, infected, processed and analyzed independently, showing mean ± SEM, with individual symbols depicting data from all culture wells. qRT-PCR samples were run in technical duplicates, with mean value plotted for each sample. Data **(D-I)** are from a single experiment in which cell subsets were isolated from a single sample containing cells pooled together from four MHV68-infected mice. Data depict 11,396 LANA.βlac-cells and 9,537 LANA.βlac+ cells. Statistical analyses **(A-C)** were performed using Mann-Whitney test, \*\*\**p* <0.001, \*\*\*\**p* <0.0001 or **(D-I)** adjusted Wilcoxon rank sum test p-values, with all statistical significance for scRNAseq comparisons indicated by †, p < 2.1x10^-56^. Supporting data in Figure S1.

### IL-4 and IFN**γ** reciprocally regulate MHV68 lytic transcription in macrophages in a time-dependent manner

To investigate how cytokines regulate the balance of MHV68 infection outcomes in macrophages, we quantified the impact of exogenous IL-4 and IFNγ on the frequency of cells initiating lytic viral gene expression, treating J774 cells with either IL-4 or IFNγ at various times pre- or post-infection (**Fig 2A**). To monitor lytic gene expression, cells were infected with MHV68.HygroGFP, with GFP expression serving as a reporter of lytic transcription^12,34^. As expected, vehicle treatment was characterized by a low (2-8%) frequency of GFP+ cells (e.g. **Fig 2B**), consistent with our previous findings^12^. IL-4 treatment resulted in a time-dependent enhancement of GFP+ cells, with maximal induction occurring with 16 hour pre-treatment (-16 hpi) (as in^17^), modest induction following 1 hour pre-treatment (-1 hpi), with little to no enhancement upon IL-4 administration at either 1 or 24 hpi (**Fig 2B-D**). Conversely, exogenous IFNγ robustly decreased the frequency of GFP+ cells, with maximal repression observed with 16 hour pre-treatment, pronounced repression with treatment either 1 hour pre- or post-infection treatment, and no discernible impact when administered at 24 hpi (**Fig 2E-F**). These data demonstrate the reciprocal regulation of MHV68 lytic replication by IL-4 and IFNγ and how these effects are impacted by time of treatment relative to de novo infection.

**Figure 2.**
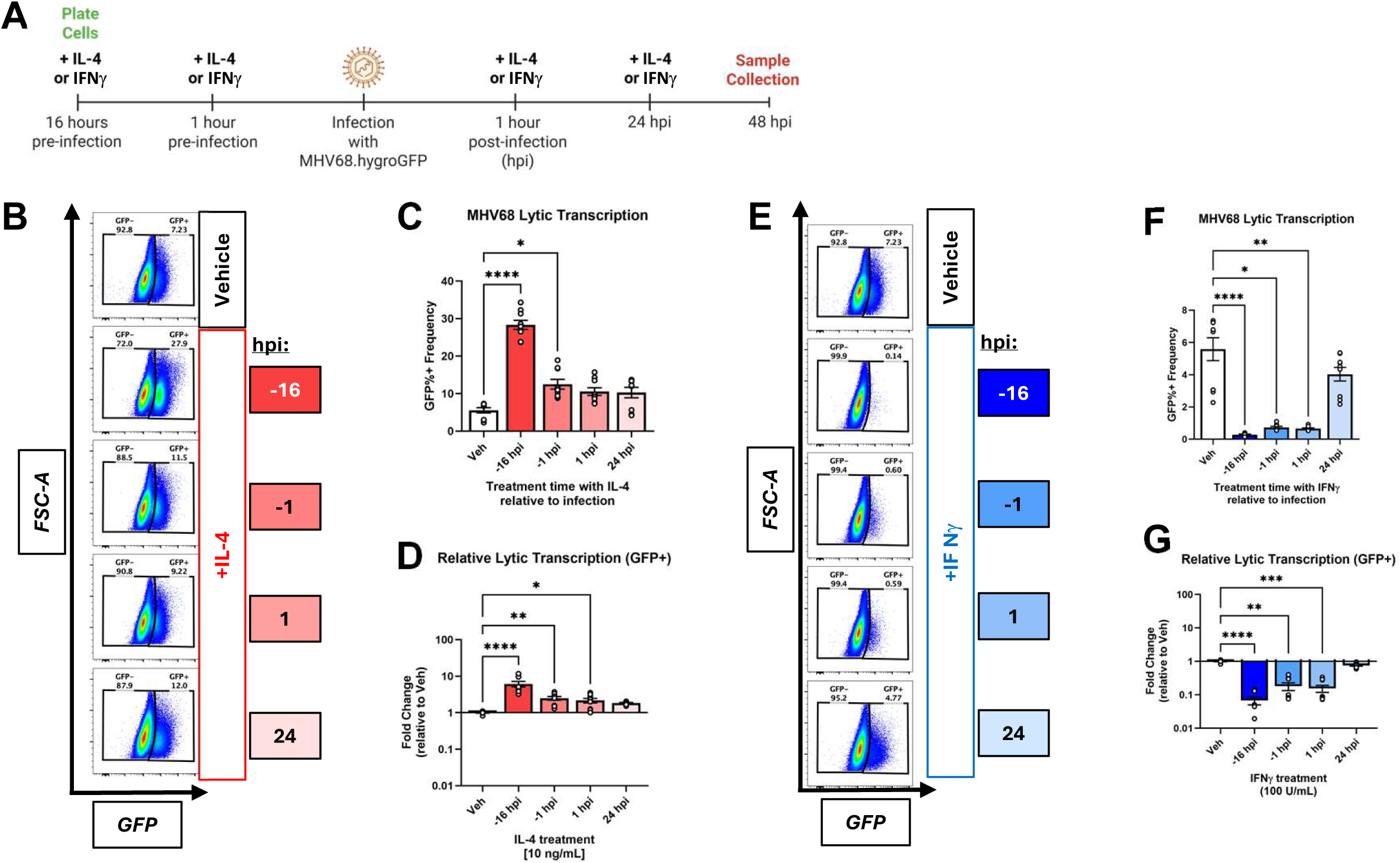
IL-4 and IFNγ reciprocally regulate MHV68 lytic transcription in macrophages in a time-dependent manner. Analysis of viral lytic cycle gene expression in MHV68-infected J774 cells subjected to cytokine treatment at various times relative to infection. **(A)** Experimental workflow, where J774 macrophages were infected with MHV68.HygroGFP (MOI=1 PFU/cell, harvested at 48 hpi), treated with either **(B-D)** IL-4 or **(E-G)** IFNγ at various times before or after infection, with the frequency of GFP+ events quantified by flow cytometry. **(B, E)** Representative flow cytometry plots quantifying the frequency of GFP+ cells (defined in the right gate) among single, viable cells, with gating strategy depicted in **Fig S2A**. **(C-D, F-G)** Quantification of the frequency of MHV68-infected J774 cells with lytic gene expression in **(C-D)** IL-4 or **(F-G)** IFNγ treated samples, as defined in panels **B, E**. Plots depict **(C, F)** the frequency of GFP+ cells or **(D, G)** fold change in the frequency of GFP+ cells relative to vehicle controls (defined as 1). For each timepoint, color shade of legend is matched between respective panels to facilitate sample cross-comparison. Data are from three independent experiments, with each experiment done in three parallel culture wells plated from a single culture, infected, processed and analyzed independently, showing mean ± SEM, with individual symbols depicting data from all culture wells. Statistical analyses for all graphs were performed using a non-parametric one-way ANOVA (Kruskal-Wallis test) comparing the mean rank of each treatment time point to vehicle treated controls. Statistical significance is indicated as \**p* <0.05, \*\**p* <0.01, \*\*\**p* <0.001, \*\*\*\**p* <0.0001. Representative flow cytometry plots for Vehicle in panels B and E are the same sample, as it served as a common control for both treatments. Supporting data in **Figure S2A**.

### Effect of cytokines and ruxolitinib, a JAK/STAT inhibitor, on virus and host gene expression, and virus replication in MHV68-infected myeloid cells

IL-4 and IFNγ binding to their respective receptors result in JAK/STAT signaling to effect gene expression^15,35^, with evidence that the promoter of the viral immediate early transactivator, *Rta*, can be regulated by both IL-4 induced STAT6 and IFNγ induced STAT1^17,26,35^. We next tested the effect of combinatorial cytokine treatment, and JAK/STAT inhibition, on infection outcome. As expected, exogenous addition of IL-4 or IFNγ enhanced or repressed the frequency of cells initiating lytic gene expression, with maximal effects observed with 16 hour pre-treatment (-16 hpi, **Fig 3A**). Parallel studies using IFNβ, another STAT1-activating cytokine, demonstrated time-dependent repression of lytic gene expression (**Fig S3A**) with a distinct anti-proliferative effect (not shown). When we tested the outcome of combinatorial cytokine treatment, cultures treated with IL-4 and IFNγ (IL-4 + IFNγ), or IL-4 and IFNβ, had a pronounced decrease in lytic infection, reminiscent of IFNγ or IFNβ treatment alone (**Fig 3A**, **Fig S3A**), consistent with a previous report^16^. In striking contrast, pre- or post-infection treatment with Ruxolitinib (RUX), a potent JAK1/JAK2 inhibitor that limits JAK/STAT signaling and is in clinical use for the treatment of certain inflammatory diseases^36^, significantly increased the frequency of cells initiating lytic replication when administered alone, or when co-administered with either IL-4 or IFNγ, irrespective of treatment time (**Fig 3A**, **Fig S3B**).

**Figure 3.**
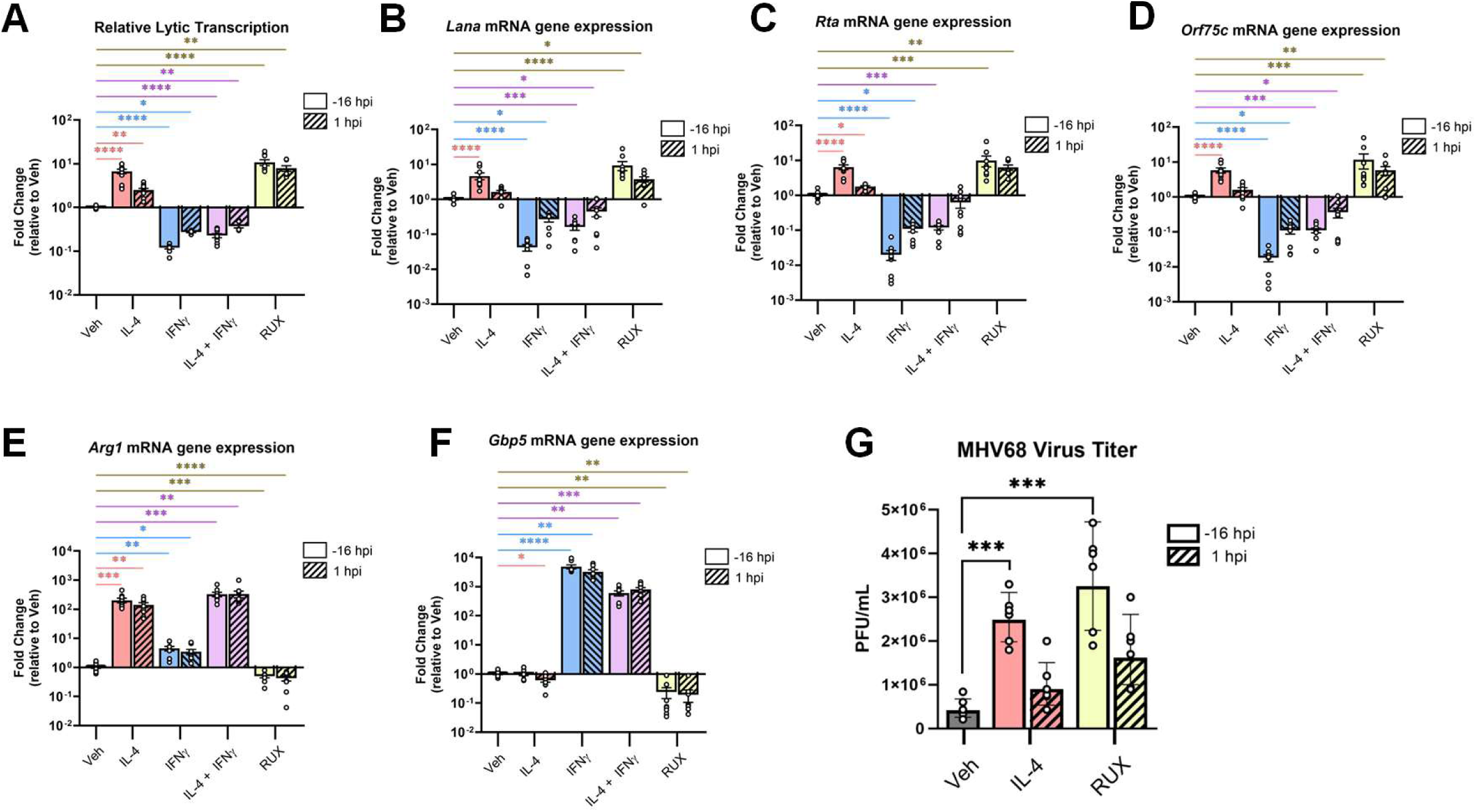
Effect of cytokines and ruxolitinib, a JAK/STAT inhibitor, on virus and host gene expression, and virus replication in MHV68-infected myeloid cells. Analysis of viral and host gene expression, and virus replication, in MHV68.HygroGFP-infected J774 myeloid cells (MOI=1 PFU/cell, harvested at 48 hpi) treated with the indicated treatment before (-16 hpi, open bars) or after (1 hpi, diagonal bars) infection. **(A)** Flow cytometry-defined frequency of GFP+ cells among single, viable cells, with gating strategy depicted in **Fig S2A. (B-F)** RT-qPCR based measurement of viral (**B**, *Lana* encoded by *Orf73*; C, *Rta*, encoded by *Orf50,* **D**, *Orf75c*) and cellular (E, *Arg1*; F, *Gbp5*) RNAs, with plots depicting fold change relative to vehicle-treated controls (set at 1), with cells treated before (-16 hpi, open bars) or after (1 hpi, diagonal bars) infection. **(G)** Infectious virus production quantified by plaque assay in cells treated with IL-4 or RUX, treated before (-16 hpi) or after (1 hpi) infection. Data are from two **(G)** to three **(A-F)** independent experiments with each experiment done in three parallel culture wells plated from a single culture, infected, processed and analyzed independently, showing mean ± SEM, with individual symbols depicting data from all culture wells. qRT-PCR samples were run in technical duplicates, with mean value plotted for each sample. Statistical analysis was done using one-way ANOVA (non-parametric, Kruskal-Wallis with Dunn’s multiple comparisons), comparing vehicle to each treatment type in individual statistical tests. Statistical significance is indicated as \**p* <0.05, \*\**p* <0.01, \*\*\**p* <0.001, \*\*\*\**p* <0.0001. Supporting data in Figure S2A, Figure S3.

Recent studies identified the epigenetic reader SP140 as a modifier of macrophage identity that regulates topoisomerase activity^37^ and interferon production^38^, which limits MHV68 lytic replication in primary macrophages^38^. To test this in our model, MHV68-infected cells were treated with an SP140 inhibitor, GSK761^39^; this treatment did not consistently alter the outcome of infection (**Fig S3C**). In contrast, combined inhibition of topoisomerase 1 and 2 with etoposide and topotecan resulted in an increased frequency of cells with lytic gene expression, but also resulted in high levels of cell death (**Fig S3C**, data not shown).

To determine the consequence of IL-4, IFNγ, RUX, and combination treatments on virus and host gene expression, we measured a panel of viral and host genes in MHV68-infected cultures. Expression of three viral transcripts (*Lana, Rta, Orf75c*) correlated with the frequency of cells initiating lytic replication (**Fig 3A**), with expression of each of these genes increased by IL-4 and RUX and decreased by IFNγ and combined treatment with IL-4 and IFNγ (**Fig 3B-D**). Expression of the IL-4 inducible gene *Arg1* was induced in both IL-4 and IL-4 and IFNγ co-treated cultures (**Fig 3E**), indicating that IL-4 signaling was not blocked upon IFNγ co-administration. Conversely, *Gbp5*, an IFN-inducible gene was prominently induced in IFNγ and combined IL-4 and IFNγ cultures, indicating that IFNγ signaling was not blocked in IL-4 and IFNγ co-administered cultures (**Fig 3F**). *Gbp5* was also decreased in RUX-treated samples, suggesting that RUX could limit basal IFN signaling in these cultures (**Fig 3F**). Finally, we assessed the impact of IL-4 or RUX on lytic replication, quantifying infectious virus production. Treatment with either IL-4 or RUX was associated with an increase in infectious virus production, an indicator of enhanced lytic replication (**Fig 3G**). These data indicate that both IL-4 treatment and JAK1/2 inhibition by ruxolitinib are associated with increased lytic replication in macrophages.

### Viral DNA replication in J774 macrophages is reciprocally regulated by IL-4 and IFN**γ** during acute in vitro infection without significant induction of viral genomic DNA methylation

Given the importance of DNA methylation as a regulator of viral gene expression and latency^29,31,40–42^, we next tested whether conditions of limited lytic replication in macrophages were associated with increased viral genomic DNA methylation. MHV68-infected J774 cells (as in **Fig 2**) were subjected to a range of conditions including baseline conditions (characterized by a low frequency of cells initiating lytic replication), conditions with enhanced lytic replication (IL-4, RUX) and conditions with further restricted lytic replication (IFNγ). Consistent with data quantifying the frequency of cells with lytic gene expression (**Fig 3, Fig S3**), baseline cultures were characterized by low viral DNA loads, with increased viral DNA loads in IL-4 and RUX-treated cultures, and decreased viral DNA loads in IFNγ-treated cultures (**Fig 4A**). In parallel, we quantified the extent of MHV68 genome methylation using a multiplex-PCR platform (iPLEX^41,42^), reading out n=73 CpG sites distributed across the viral genome. We observed no significant DNA methylation across any treatment type or time point, in contrast to the MHV68 latently infected A20.HE2.1 B cell line^34^ (**Fig 4B**). When we quantified viral DNA load and methylation in MHV68-infected J774 cells or 3T12 fibroblasts, a cell type characterized by robust lytic replication, viral DNA load was increased as a function of time (**Fig 4C-D**), with negligible viral genome methylation in either cell type at these acute timepoints (**Fig 4E-F**). These data suggest that viral DNA methylation is not a prominent mechanism involved in limiting de novo MHV68 macrophage infection outcome.

**Figure 4.**
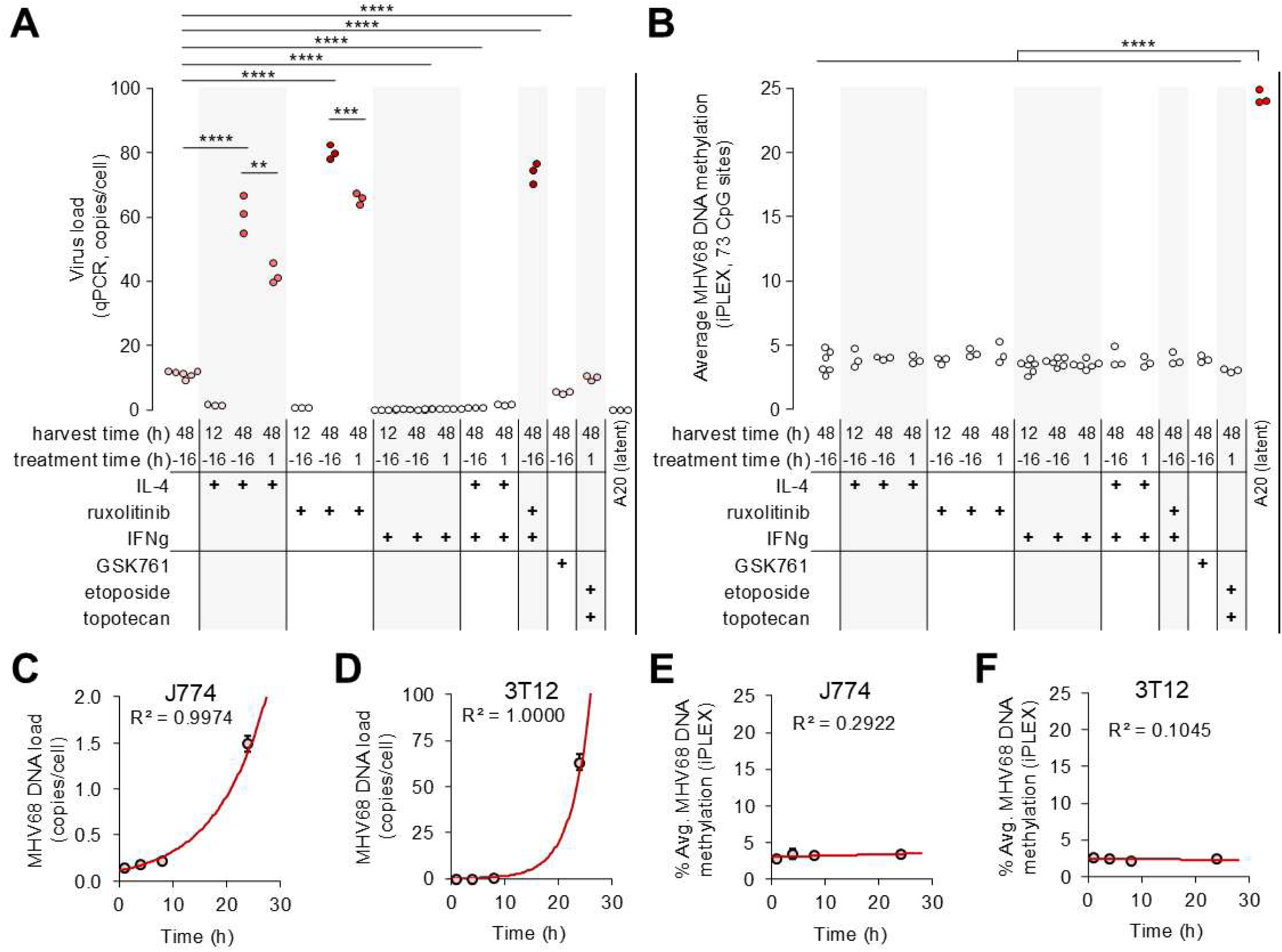
Viral DNA replication in J774 macrophages is reciprocally regulated by IL-4 and IFNγ during acute in vitro infection without significant changes to viral genomic DNA methylation. Analysis of viral DNA and viral genomic DNA methylation following WT MHV68 infection (MOI=1 PFU/cell) of J774 macrophages or 3T12 fibroblasts. Analysis also included A20.HE2.1 cells, an MHV68-infected, latent B cell line, “A20 (latent)”. **(A-B)** Quantitation of **(A)** viral DNA by qPCR, measuring viral DNA copies per cell or **(B)** percent average viral genomic DNA Methylation quantified by iPLEX across 73 CpG sites, in MHV68-infected J774 cells subjected to the indicated conditions and harvested at the indicated time. **(C-F)** Quantification of **(C-D)** viral DNA by qPCR, measuring viral DNA copies per cell or **(E-F)** percent average viral genomic DNA methylation quantified by iPLEX in either **(C, E)** J774 macrophages or **(D, F)** 3T12 fibroblasts over time. One independent experiment was performed with three biological replicate samples. Each biological replicate was run in technical duplicates. Statistical significance was determined by unpaired, two-tailed t-test where \**p* <0.05, \*\**p* <0.01, \*\*\**p* <0.001, \*\*\*\**p* <0.0001.

### IFN**γ** pretreatment of macrophages reduces lytic infection and transcription of a virus that constitutively expresses the Replication and Transcription Activator (RTA)

Previous data have implicated transcriptional regulation of RTA, encoded by the *Orf50* gene of MHV68, as a major point of cytokine regulation^16,26^. To gain insight into the molecular mechanistic effects of IFNγ, we tested the impact of IFNγ on the outcome of infection, using a recombinant MHV68 (MHV68.C-RTA) that undergoes constitutive lytic replication. In this virus, in addition to the native *Orf50* gene, an additional RTA gene was inserted into the left end of the genome, with constitutive, high-level expression driven by the HCMV IE promoter^32^. To assess lytic gene expression, cultures were analyzed for the frequency of cells expressing proteins associated with MHV68 lytic replication: i) the viral regulator of complement activation (vRCA, encoded by *Orf4*), and ii) γH2AX, a histone phosphorylation event robustly induced by an MHV68 viral kinase^43,44^ and/or cell stress. As expected, WT MHV68-infected J774 cells had a low frequency (<1%) of cells co-expressing vRCA and γH2AX (**Fig 5A-B**), consistent with other measures of limited lytic replication in this system (**Fig 2**). In contrast, J774 cells infected with MHV68.C-RTA demonstrated a high (>50%) frequency of cells co-expressing vRCA and γH2AX (**Fig 5A-B**), indicating that RTA overexpression in this system is capable of driving lytic replication in a large percentage of cells. Notably, IFNγ pretreatment of C-RTA infected cells resulted in a dramatic decrease in the frequency of vRCA- and γH2AX-expressing cells relative to vehicle-treated C-RTA infection cultures, to more closely resemble infection outcomes with WT MHV68 (**Fig 5A-B**). Changes in the frequency of cells with lytic gene expression were mirrored by quantification of *Rta* and *Lana* transcripts, with vehicle-treated C-RTA infected cultures characterized by greatly increased expression of *Rta* and *Lana* relative to WT infected cultures (**Fig 5C-D**). While IFNγ-treated C-RTA infected cultures had modestly increased expression of *Lana* relative to WT infected cultures, *Rta* and *Lana* expression was significantly reduced relative to vehicle treated C-RTA infected cultures (**Fig 5C-D**). These data indicate that IFNγ can potently limit MHV68 lytic replication even in the context of RTA overexpression.

**Figure 5.**
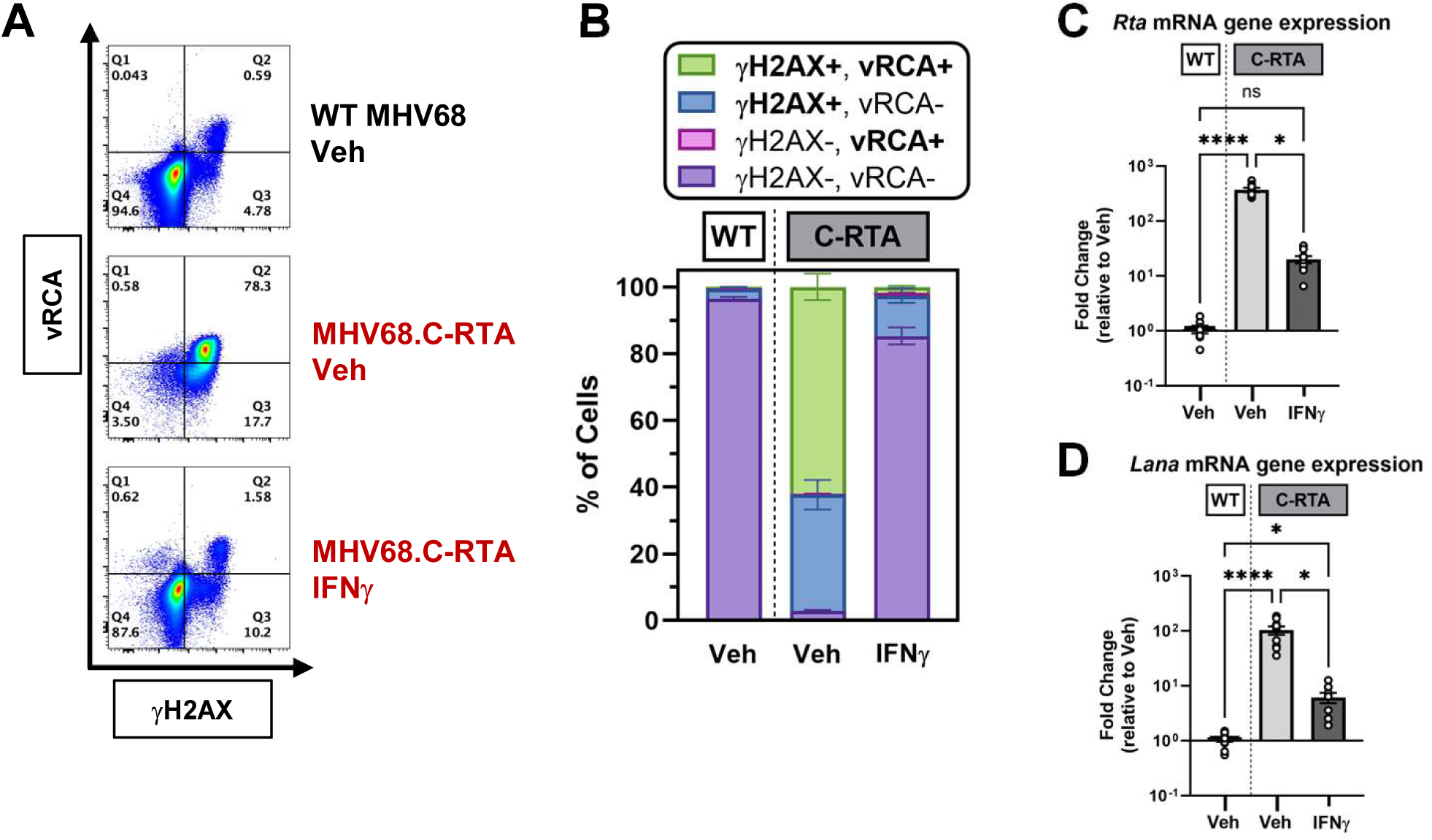
IFNγ pretreatment of macrophages reduces lytic infection and transcription of a virus that constitutively expresses the Replication and Transcription Activator (RTA). Analysis of virus infection outcomes in J774 macrophages comparing infection with WT or C-RTA MHV68 (MOI=1 PFU/cell) with or without IFNγ treatment, harvested at 48 hpi. **(A-B)** Flow cytometric analysis of proteins associated with MHV68 lytic replication (vRCA, γH2AX) in single, viable cells with gating strategy in Fig S2B. The frequency of cells characterized by vRCA and/or γH2AX expression is quantified in **(A)** representative flow cytometry plots across **(B)** all experiments. **(C-D)** RT-qPCR quantitation of MHV68 RNAs for **(C)** *Rta* encoded by *Orf50* and **(D)** *Lana* encoded by *Orf73* RNA in MHV68-infected J774 macrophages, pre-treated either with vehicle (Veh) or IFNγ at -16 hpi. Data are from three independent experiments with each experiment done in three parallel culture wells plated from a single culture, infected, processed and analyzed independently, showing mean ± SEM, with individual symbols depicting data from all culture wells. Statistical analyses for panels **C-D** were performed using a non-parametric one-way ANOVA (Kruskal-Wallis test) with Dunn’s multiple comparisons test, comparing all conditions to each other. \**p* <0.05, \*\*\*\**p* <0.0001. Supporting data in **Figure S2B**.

### IFN**γ** pretreatment of macrophages is associated with a reduced frequency of LANA expressing cells and reduced nuclear viral DNA

To determine whether IFNγ alters early steps in virus infection, we used the MHV68.LANA::βlac reporter virus (β-lactamase fusion to the immediate early viral gene, *Lana*, expressed during both latent and lytic infection^33^) to quantify the frequency of cells in which virus infection has progressed from attachment, entry, import of the viral genome into the nucleus, culminating in viral transcription. Studies compared the effect of IFNγ treatment pre- or post-infection. Vehicle-treated cultures demonstrated a high frequency (∼60%) of LANAβlac+ cells, with similar infection frequencies observed in cultures treated with IFNγ (1 hour post-infection) or treated with GSK761, the SP140 inhibitor, pre- or post-infection (**Fig 6A**). In contrast, cultures pre-treated with IFNγ for 16 hours (-16 hpi) showed a pronounced decrease in the frequency of LANAβlac+ cells, a reduction also observed in cultures co-treated with IFNγ and GSK761 (**Fig 6A**), suggesting that IFNγ pre-treatment may impede virus infection in a significant fraction of cells. To elucidate how IFNγ pre-treatment influences the progression of infection, we quantified viral DNA as a proxy for virus attachment, entry and nuclear entry as detailed in **Fig 6B**. Studies compared these stages of infection between vehicle- or IFNγ-treated cultures across a threefold series of input infection doses. Vehicle- or IFNγ-treated cultures had comparable viral DNA in assays for attachment (assessed at 1 hpi) and entry (assessed at 3 hpi) (**Fig 6C**). In contrast, IFNγ pretreated cultures demonstrated a consistent reduction in nuclear virus DNA (assessed at 5 hpi) across all infection doses tested (**Fig 6D**). This reduction was similar in magnitude to that observed with IFNγ pretreatment of LANAβlac+ cells (**Fig 6A**). These data suggest that IFNγ pretreatment either limits nuclear entry of the viral genome or decreases viral genome stability within the nucleus.

**Figure 6.**
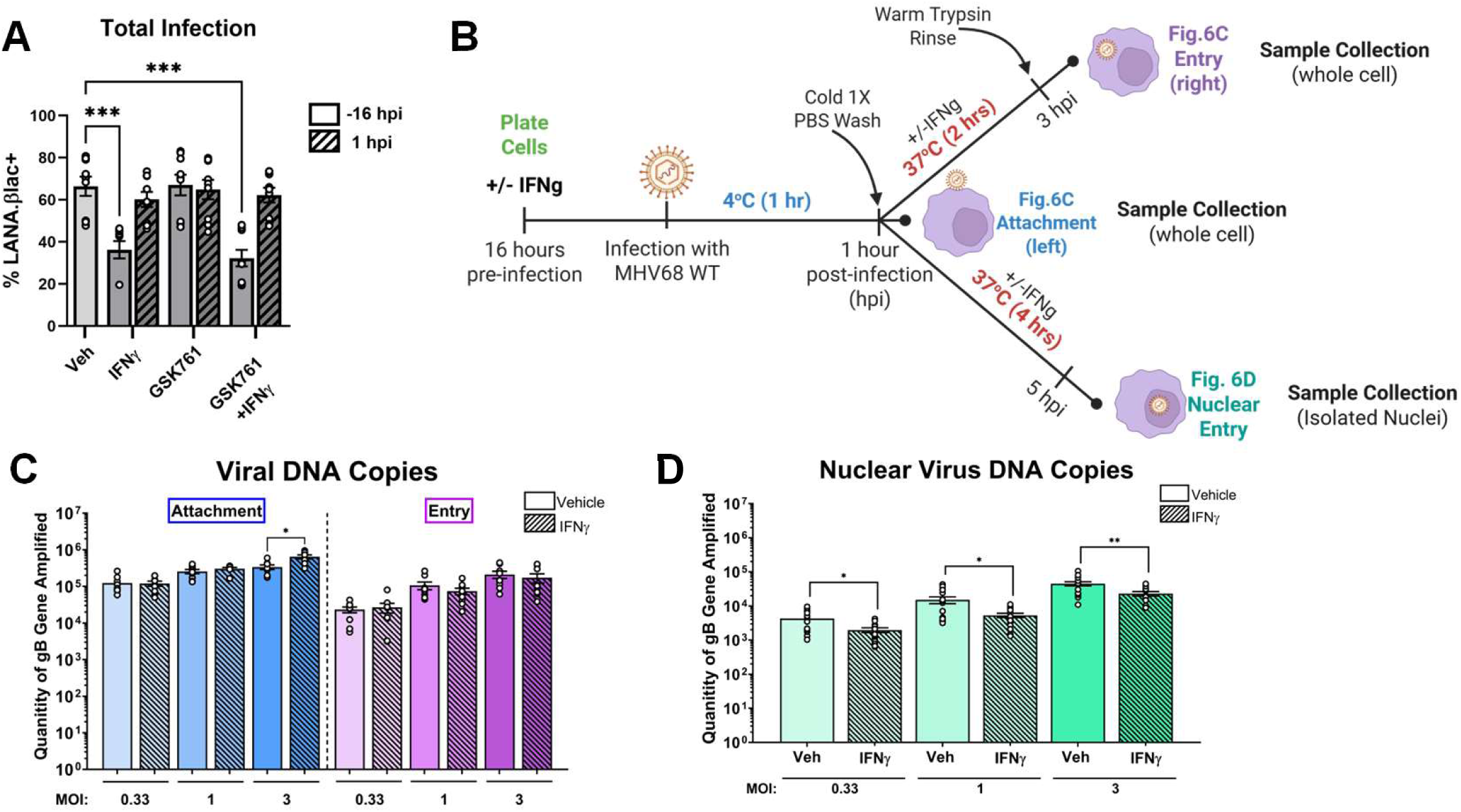
IFNγ pretreatment of macrophages is associated with a reduced frequency of LANA expressing cells and reduced nuclear viral DNA. Analysis of viral transcription and viral DNA in distinct stages of infection in MHV68-infected J774 macrophages as a function of IFNγ treatment. **(A)** Flow cytometric analysis of the frequency of LANAβlac+ cells in MHV68.LANA::βlac-infected J774 macrophages (MOI=1 PFU/cell, harvested at 48 hpi) subject to the indicated treatments, given before (-16 hpi, open bars) or after (1 hpi, diagonal bars) infection. The frequency of LANAβlac+ cells defined by β-lactamase substrate cleavage among single, viable cells with gating strategy shown in **Fig S2C**. Data shown represent the percentage of single, viable, substrate cleavage-positive cells to identify the LANAβlac expression. **(B)** Graphical representation of the experimental workflow used for data presented in panels **C-D**. **(C-D)** Viral DNA quantitation by qPCR to assess (**C**, left plot, shades of blue) virus target cell attachment, (**C**, right plot, shades of purple) viral entry into the host cell and (**D**, green) viral DNA localization within the nucleus using the experimental methods detailed in panel **B**. Cells were infected at a range of MOIs (as indicated, x axis) and pre-treated with vehicle (Veh, open bars) or IFNγ (diagonal bars). Data are from two **(D)** to three **(A-C)** independent experiments with each experiment done in three parallel culture wells plated from a single culture, infected, processed and analyzed independently, showing mean ± SEM, with individual symbols depicting data from all culture wells. PCRs were run in technical duplicates, with mean value plotted for each sample. Statistical analysis in **A** was performed using a Kruskal-Wallis test with Dunn’s multiple comparisons test. \*\*\**p* <0.005. Statistical analyses in **C-D** were performed using a Mann-Whitney test for comparisons within each individual MOI and stage of infection. \**p* <0.05, \*\**p* <0.01. Veh, Vehicle; IFNγ, interferon gamma; GSK761, an Sp140 inhibitor. Supporting data in **Figure S2C.**

### IFN**γ** treatment of macrophages reveals divergent regulation of latent and lytic gene expression

The ability of IFNγ to limit MHV68 infection with pre- or post-infection treatment (**Fig 2**), combined with the ability of IFNγ to limit the frequency of LANAβlac+ cells (**Fig 6**) strongly suggested multiple levels of cytokine-dependent control of infection. To detect the proportion of cells undergoing lytic versus total infection within the same cell population, we used reporter viruses with the β-lactamase gene either fused in-frame to the endogenous genes *Lana* (MHV68.LANA::βlac; encoding an immediate early gene expressed in all infected cells) or *Orf72* (MHV68.vCyc::βlac; encoding the early-late *vCyclin* gene expressed only in cells initiating lytic early gene transcription), to sensitively monitor endogenous viral gene expression with a common readout. To demonstrate the utility of this system, parallel cultures of J774 cells were infected with WT MHV68 (negative control), MHV68.LANA::βlac, or MHV68.vCyc::βlac, subjected to fluorescent-barcoding and quantified for the frequency of LANA+ cells, a marker of total infection, relative to vCyc+ cells, a marker of lytic infection (**Fig 7A**). As predicted, the frequency of LANA.βlac expressing cells in these cultures was significantly higher than the frequency of vCyc.βlac expressing cells (**Fig 7B-C**), consistent with multiple measures demonstrating efficient infection of these cells, with a low frequency of cells initiating lytic infection (**Fig 2**). IL-4 treatment of these cultures demonstrated a time-dependent enhancement of both lytic and total infection, as measured by increased frequencies of vCyc.βlac- and LANA.βlac-expressing cells by 48 hpi (**Fig 7D**). The higher frequency of LANA.βlac-expressing cells compared to vCyc.βlac expressing cells was further observed during primary in vivo infection of mice (**Fig 7E**).

**Figure 7.**
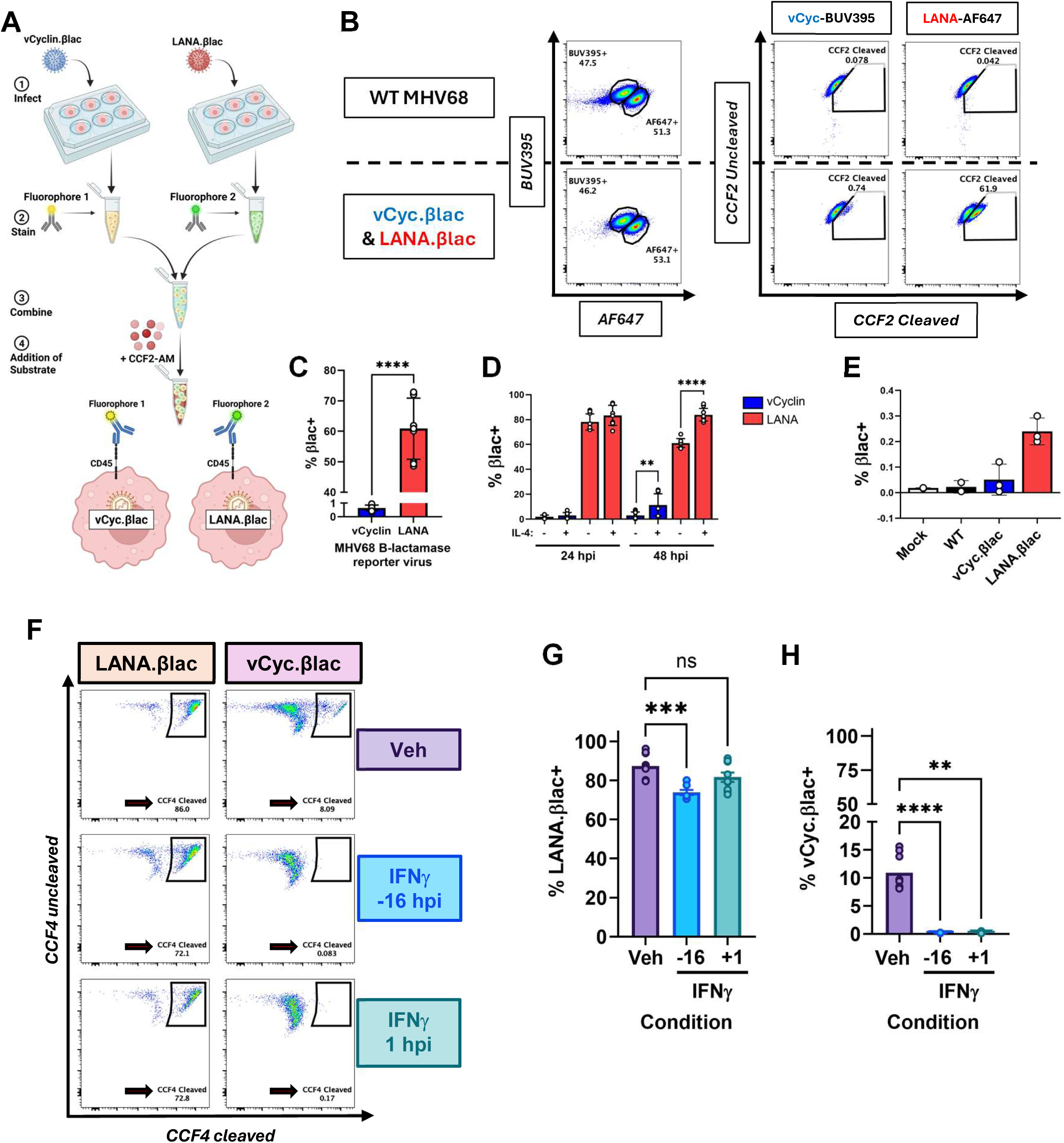
IFNγ treatment of macrophages reveals divergent regulation of latent and lytic gene expression. Analysis of LANA and vCyclin gene expression in MHV68-infected J774 macrophages and primary peritoneal macrophages as a function of IFNγ treatment. **(A)** Experimental workflow to simultaneously assess LANA and vCyclin reporter gene expression in MHV68-infected J774 cells (MOI=1 PFU/cell, harvested at 48 hpi) by flow cytometry. (1) Independent cultures were infected with MHV68.LANA::βlac or MHV68.vCyc::βlac, harvested at 48 hpi, (2) incubated with distinct fluorescently labelled antibodies (BUV395- or AF647-labelled anti-CD45 antibodies) to barcode each sample, (3) combined into a single reaction tube, (4) incubated with a fluorescent β-lactamase substrate, analyzed for the frequency of cells with substrate cleavage by flow cytometry. **(B)** Representative flow cytometry plots of single, viable cells to define fluorescently-barcoded populations (left column), with CCF2 loading and substrate cleavage quantified in either BUV395- (middle column) or AF647-labelled populations (right column). Data compare WT MHV68-infected cells (top row, without β-lactamase reporter) to define background fluorescence, with β-lactamase reporter virus-infected cells (bottom row), with viral gene expression defined based on cleaved substrate fluorescence (full gating strategy shown in **Fig S2D**)**. (C-D)** Quantitation of the frequency of vCyc.βlac+ or LANA.βlac+ cells in MHV68-infected J774 cells (MOI=1 PFU/cell) **(C)** in the absence of additional stimuli at 48 hpi, or **(D)** in the presence or absence of IL-4 at 24 or 48 hpi. **(E)** In vivo analysis of the frequency of vCyc.βlac+ or LANA.βlac+ cells in MHV68-infected BALB.IFNγ-deficient mice (1x10^6^ PFU, i.p. injection, harvested at 16 hpi), quantifying the frequency of βlac+ cells among single, viable, CD45+ peritoneal cells, with individual symbols indicating individual mice. **(F-H)** Ex vivo analysis of the frequency of vCyc.βlac+ or LANA.βlac+ cells among primary peritoneal cells infected ex vivo (MOI=1 PFU/cell, harvested at 48 hpi), comparing vehicle (Veh) or IFNγ treatment before (-16 hpi) or after (1 hpi) infection, analyzing reporter gene expression in single, viable, peritoneal macrophages (defined by expression of CD11b and F4/80) (full gating strategy in **Fig S2E**). Data are from (**B-D, F-H**) three independent experiments with each experiment done in three parallel culture wells plated from a single culture, infected, processed and analyzed independently or (**E**) 1 experiment. Data depict mean ± SEM with individual symbols indicating **(B-D, F-H)** individual culture wells or **(E)** individual mice. Statistical analyses for panel **C-D** were performed using a Mann-Whitney test, with statistical analysis of panel **D** comparing the presence or absence of IL-4 treatment for each reporter virus at each analysis time point. Statistical analysis for panels **F-G** were performed using a non-parametric one-way ANOVA (Kruskal-Wallis test) comparing the mean rank of each treatment and time point to vehicle treated controls. Statistical significance is indicated as \*\**p* <0.01, \*\*\**p* <0.001, \*\*\*\**p* <0.0001. Supporting data in **Figure S2D-E.**

To test the impact of IFNγ on lytic and total infection in primary macrophages, we collected primary peritoneal cells from naïve WT C57BL/6 mice and infected them ex vivo, comparing vehicle-treated or IFNγ treated cultures, with IFNγ added before (-16 hours) or after (1 hour) infection. Vehicle-treated peritoneal macrophages were characterized by a high frequency (>80%) of LANA.βlac+ cells with a low frequency (∼10%) of vCyc.βlac+ cells (**Fig 7F-H**). In cultures pretreated with IFNγ, peritoneal cells had a decreased frequency of LANA.βlac+ cells, with a more pronounced decrease in vCyc.βlac+ cells; in contrast, cultures where IFNγ was added 1-hour post-infection had comparable frequencies of LANA.βlac+ to vehicle-treated cultures, with a pronounced decrease in vCyc.βlac+ cells (**Fig 7F-H**). These data strongly suggest that IFNγ is capable of impacting both the total frequency of infection and the extent of lytic infection, depending on when macrophages are treated with IFNγ.

## DISCUSSION

The outcome of γHV infection is intimately regulated by the dynamic balance between lytic infection and latency, a process substantially shaped by the host immune response. In particular, cytokines play a major role in immune regulation of virus infection, especially in myeloid cells, with IFNγ and IL-4 identified as key determinants that either limit or promote virus replication^16–19^. Further, disruption of these pathways, in conditions of immune-deficiency or co-infection, can profoundly alter the outcome of infection at the organismal level^16,21–24,45^.

Previous studies using the MHV68 system have identified myeloid cells as a key link between lytic replication in epithelial cells and latent infection in B cells^10^, with macrophages both a prominent early target of infection after multiple routes of infection^8,46^ and a potential latent reservoir^9,11^. Our recent studies further identified that macrophages are readily infected by MHV68, resulting in a substantial fraction of cells characterized by limited virus transcription, a restricted, latent-like state of infection, with rare cells initiating lytic replication. Here, we expanded upon these findings and sought to elucidate how two well-documented regulators of MHV68 macrophage infection, IL-4 and IFNγ, influenced infection outcomes.

Our data clearly demonstrates that IL-4 and IFNγ have the ability to alter the frequency of cells initiating lytic replication with distinct temporal kinetics. Whereas IL-4 pretreatment profoundly enhances the frequency of cells initiating lytic replication, IL-4 administration after de novo infection had minimal impact on lytic replication, despite the fact that IL-4 induced *Arg1*, a canonical IL-4 transcriptional target. This finding suggests IL-4 induction of STAT6, a rapid post-translational response, to transactivate the *Rta* promoter^16^ may not be the sole mechanism by which IL-4 promotes lytic infection. In contrast, we find that IFNγ treatment before or shortly after infection is able to impair lytic gene expression. Our data suggest a two-stage model by which IFNγ can limit lytic infection. First, when cells are exposed to IFNγ after infection, it is likely that IFNγ represses lytic replication through a rapid post-translational mechanism, potentially through STAT1 activation, nuclear translocation and repression of *Rta* transcription, a mechanism proposed by Goodwin et al^26^. Notably, IFNγ treatment after infection does not impact the frequency of LANA-expressing cells, strongly suggesting that IFNγ selectively interferes with lytic gene expression. Second, when cells are exposed to IFNγ before infection, we find a decreased frequency of LANA-expressing cells, reduced nuclear viral DNA, as well as profound restriction of lytic infection. We posit that IFNγ treatment before infection elicits a second level of control, inducing effector mechanism(s) that limit virus translocation to the nucleus or reduce virus genome stability in the nucleus. Additionally, IFNγ pretreatment is able to restrict the frequency of cells initiating lytic replication, a mechanism that resembles that observed with IFNγ exposure after infection.

While the molecular mechanisms by which IFNγ limits MHV68 infection and lytic replication remain to be determined, our data suggest additional complexity. For example, the native *Rta* promoter has been shown to be inhibited by IFNγ in a dose-dependent, STAT1-dependent manner^26^. One potential complexity to this model is our data showing that IFNγ can potently reduce lytic cycle progression following infection with the C-RTA virus, in which RTA is expressed from both its native locus as well as from a second locus under control of the HCMV IE promoter^32^. While these studies used IFNγ pre-treatment rather than post-treatment, the ability of IFNγ to restrict RTA expression and lytic cycle progression even when RTA is transcribed from a strong, heterologous promoter suggests that IFNγ may be able to regulate infection downstream of *Rta* transcription. While our current data do not support an immediate role for viral genomic DNA methylation for restricted macrophage infection under basal or IFNγ treated conditions during acute infection, future studies will be required to investigate more proximal mechanisms of epigenetic silencing (e.g. histone modifications^27,30,40^). In addition, it stands to reason that DNA methylation may only be involved in viral gene silencing at later timepoints after infection, as evidenced for the related human herpesvirus KSHV^47^.

Our studies also emphasized the dominant role of basal JAK/STAT signaling as an inhibitor of lytic infection in macrophages, a finding revealed by use of the JAK1/2 inhibitor, ruxolitinib. Indeed, treatment with ruxolitinib (either pre- or post-infection) robustly increased the frequency of cells initiating lytic replication, ultimately culminating in increased lytic replication and infectious virus production. Though the in vivo relevance of this remains to be experimentally tested, it is notable that ruxolitinib is now used in patients to treat inflammatory diseases, including topical application for eczema and systemic administration for graft versus host disease^36^. The impact of ruxolitinib, or other JAK inhibitors, may have complex impacts on the control of the human γHVs, including EBV^48,49^, depending on the clinical context.

In total, our findings provide new insights into how IL-4, IFNγ and JAK/STAT signaling regulate the outcomes of γHV infection in myeloid cells. These studies emphasize the distinct impacts that these pathways have, and how exposure before or after infection can profoundly alter their efficacy and infection outcomes. Our studies further emphasize the importance of studying these processes with single cell level resolution when possible, as these effects are not all or none, instead changing the frequencies of cells that achieve highly divergent outcomes. Because macrophages are an early target of γHV infection and have been shown to facilitate B cell infection, IFNγ mediated regulation at multiple stages of infection may have additional consequences on the magnitude of latency establishment during primary infection. We anticipate that these findings may have broader relevance to the human γHVs, EBV and KSHV, potentially regulating infection outcomes in myeloid cells and the dynamic control of latency and reactivation.

### Limitations of Study

This study has important limitations. First, our studies relied heavily on the J774 macrophage cell line, and while the central findings were corroborated in primary peritoneal macrophages, macrophages exhibit profound phenotypic and functional diversity, heavily informed by tissue-specific cues^50,51^. Conservatively, we can conclude that IFNγ has the potential to limit primary de novo infection of primary peritoneal macrophages, a prominent target of early MHV68 infection^8^. Whether these findings extend to other macrophage subsets infected by MHV68 (e.g. alveolar macrophages or subcapsular sinus macrophages^46,52^) and how these findings translate in vivo remain unknown. Second, these studies focused on a single viral infection dose and single treatment doses, demonstrating a key temporal component for cytokine effects, adding to the body of literature investigating how IFNγ constrains γHV infection. It is very likely that distinct mechanisms are elicited with different infection doses and cytokine concentrations, an area for future investigation. At this time, the molecular mechanism(s) by which IFNγ and JAK/STAT signaling contain different infection stages remain unidentified. Though our studies did not find evidence for significant viral genomic DNA methylation during de novo infection in vitro, we anticipate that this mechanism will be highly relevant during latent infection of macrophages and B cells in vivo^31^. Our studies investigating the contribution of SP140 and topoisomerases as regulators of macrophage infection rely on pharmacologic inhibitors and may require genetic corroboration to provide definitive answers.

## RESOURCE AVAILABILITY

### Lead contact

Further information and requests for resources and reagents should be directed to and will be fulfilled by the lead contact, Linda F. van Dyk.

### Materials availability

This study did not generate new unique reagents

### Data and code availability

• Raw and processed single cell RNA-Seq data are available through NCBI GEO (GSE345343), publicly available as of the date of publication.

• This article does not report original code.

• Any additional information required to reanalyze the data reported in this paper is available from the lead contact upon request.

## ACKNOWLEDGEMENTS

We thank Christine Childs and Kristina Terrell of the University of Colorado Cancer Center Flow Cytometry Shared Resource for training and assistance with flow cytometry and Dr. Tonya Brunetti in the Bioinformatics and Biostatistics Shared Resource for assistance with scRNA-seq data deposition. We thank Dr. Craig Forrest for generously sharing the MHV68.HygroGFP recombinant virus, and Dr. Ren Sun for generously sharing the MHV68.C-RTA recombinant virus. This study was supported by the National Institutes of Health grant RO1 AI157201 awarded to LvD and ETC, Molecular Biology T32 pre-doctoral training grant (NIH5T32GM136444) awarded to REK, an American Cancer Society Center for Innovation in Cancer Research Training grant (POST-BACC-23-1158368-01-DPBACC) awarded to P.G., with additional support by the National Institutes of Health Cancer Center Support Grant (P30CA046934), including the Bioinformatics and Biostatistics Shared Resource (RRID: SCR_021983), Flow Cytometry Shared Resource (RRID: SCR_022035) and Genomics Shared Resource (RRID: SCR_021894).

## AUTHOR CONTRIBUTIONS

Conceptualization: R.E.K., E.T.C., L.F.v.D.;

Methodology: C.W., R.A.B.; D.G.O.;

Validation: R.E.K., G.V., P.G., C.W., E.M.M.;

Formal Analysis: A.G., C.W., E.T.C

Investigation: R.E.K, G.V., P.G., C.W., E.M.M.;

Data Curation: R.E.K., A.G., E.T.C.;

Writing – original draft: R.E.K., E.T.C., L.F.v.D.;

Writing – review & editing: R.E.K, E.T.C., L.F.v.D.;

Visualization: R.E.K., C.W., A.G., E.T.C., L.F.v.D.;

Supervision: E.T.C., L.F.v.D.;

Project administration: R.E.K., E.T.C., L.F.v.D.;

Funding acquisition: E.T.C., L.F.v.D.

## DECLARATION OF GENERATIVE AI AND AI-ASSISTED TECHNOLOGIES IN THE WRITING PROCESS

Generative AI was not used in preparation of this manuscript.

## STAR METHODS

### Key resources table

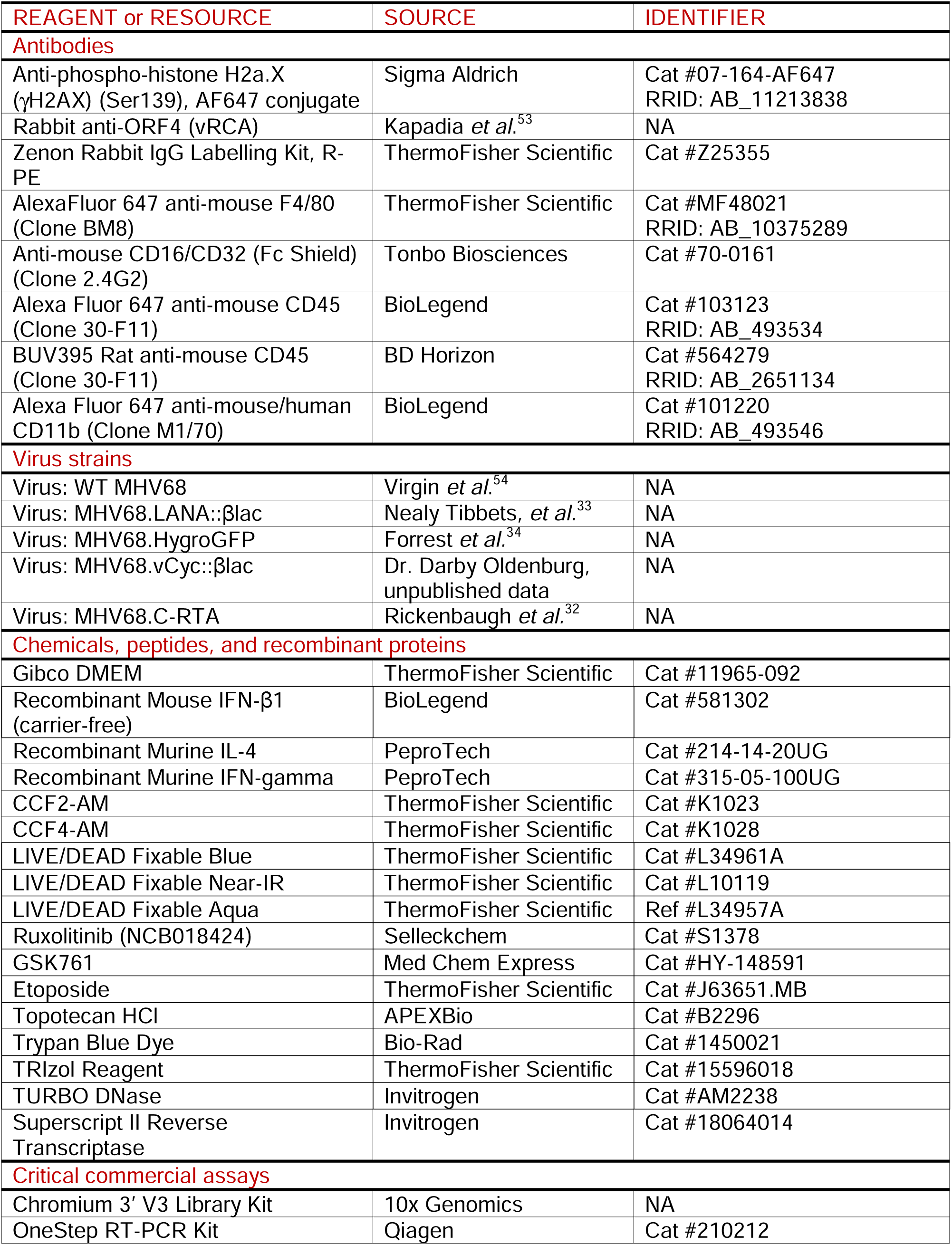
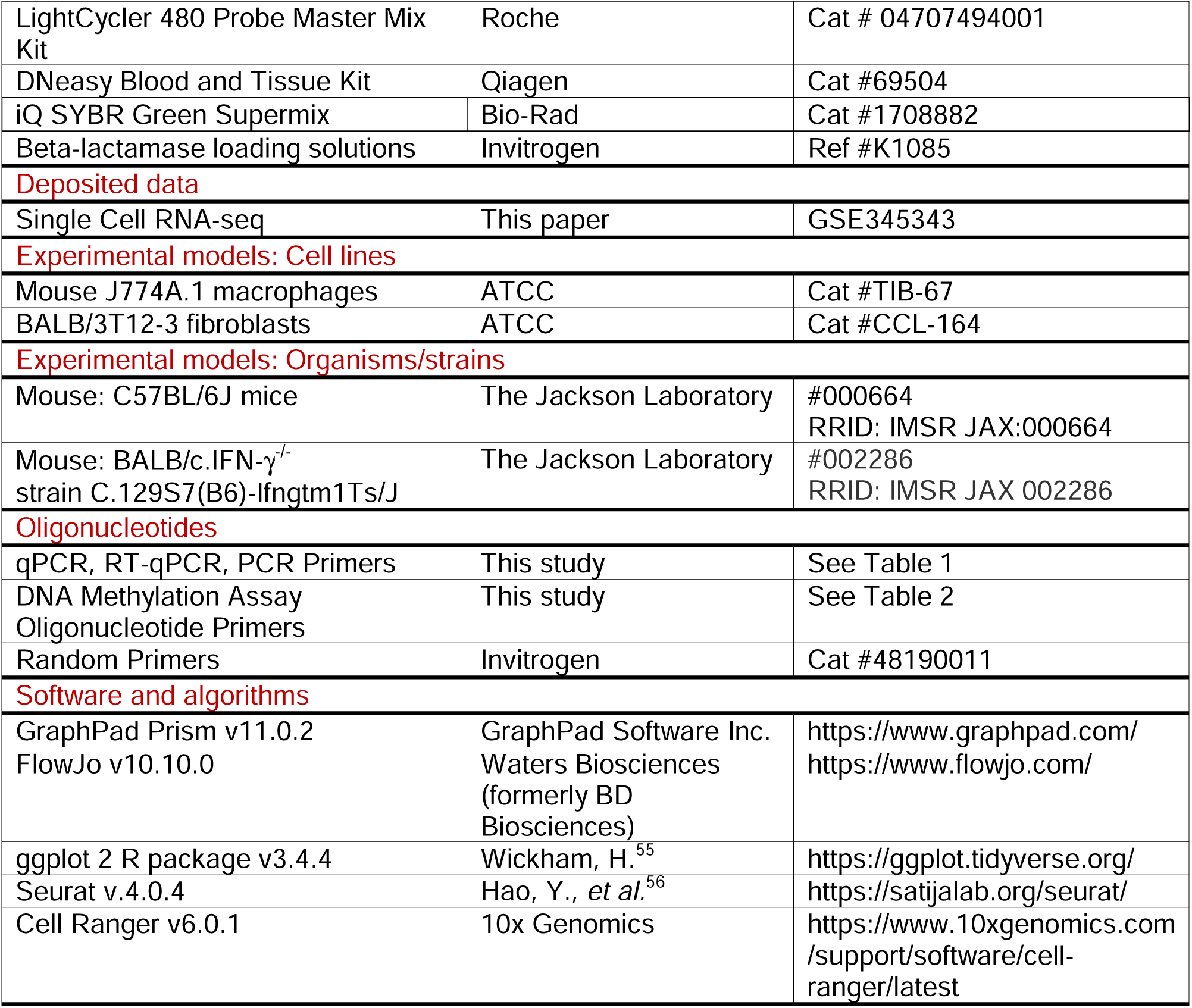

## METHODS

### Mice

WT C57BL/6J female mice (stock #000664) were obtained from the Jackson Laboratory (Bar Harbor, ME). BALB/c.IFNγ^-/-^ mice, originally obtained from the Jackson Laboratory [strain C.129S7(B6)-IFNγtm1Ts/J, stock #002286] were bred in-house at the University of Colorado Denver Anschutz Medical Campus in accordance with university regulations. Mice were infected between 8 and 25 weeks of age. Infected mice were housed in an animal biosafety level 2 facility in accordance with University regulations. Mice were infected intraperitoneally with WT MHV68 or indicated reporter viruses at a dose of 1x10^6^ PFU in a 500 µL total volume of complete DMEM, harvested at the indicated times.

### Cells and Viruses

Mouse J774A.1 macrophages (subsequently referred to as “J774”; ATCC TIB-67) were cultured in complete DMEM (cDMEM): Dulbecco’s modified Eagle medium (DMEM; Life Technologies) supplemented with 10% fetal bovine serum (FBS; Atlanta Biologicals), 2 mM L-glutamine, 10 U/mL penicillin, and 10 μg/mL streptomycin sulfate. Mouse 3T12 fibroblast cells (ATCC CCL-164) were cultured in cDMEM with 5% FBS. All cells were cultured at 37°C with 5% CO_2_. A20.HE2.1 cells, an MHV68-infected, latent B cell line was grown as described^34^. Wild-type (WT) MHV68 (WUMS), MHV68.HygroGFP, MHV68.LANA::βlac, MHV68.vCyclin::βlac, and MHV68.C-RTA viruses (**Fig S4**) were grown and prepared as previously described^33,34,57^, with virus titers quantified by at least three independent plaque assays. MHV68.vCyc::βlac was created through BAC recombineering, introducing an in-frame, C-terminal fusion with the βlac gene downstream of the endogenous vCyc (ORF72) gene (kindly provided by Dr. Darby Oldenburg, unpublished data). All recombinant viruses used in these studies had the BAC origin of replication removed prior to viral stock generation. All recombinant viruses used in this study are capable of robust, lytic replication in fibroblasts, replicating to high-titers of infectious virus with kinetics comparable to WT MHV68 (data not shown), with the exception of MHV68.C-RTA which is known to have accelerated lytic replication kinetics^32–34,57,58^.

In vitro and ex vivo infections were done using a multiplicity of infection (MOI) of 1 PFU/cell, based on live cell counts using a TC20 Automated Cell Counter (Bio-Rad) with Trypan Blue dye. Infections were done in 12-well or 6-well plate format, with cells incubated with virus at 37°C with 5% CO_2_ for 1 hour, in a volume of 150 μL (12-well) or 300 μL (6-well), rocking every 15 minutes. Viral inoculum was removed and replaced with 1mL cDMEM at 1 hour (unless otherwise indicated). For cytokine and pharmaceutical treatments, media containing treatment was added to cells at indicated times pre- or post-infection, with cells harvested by cell scraping (for DNA isolation or flow cytometry), or incubation in TRIzol Reagent (for RNA isolation). Measures of infection included: viral DNA quantitation by quantitative real-time PCR, viral and host mRNA quantitation by RT-qPCR (PCR primers, **Table 1**), and infectious virus quantitation by plaque assay (as in^12^, detailed in **Supplemental Methods**).

**Table 1:**
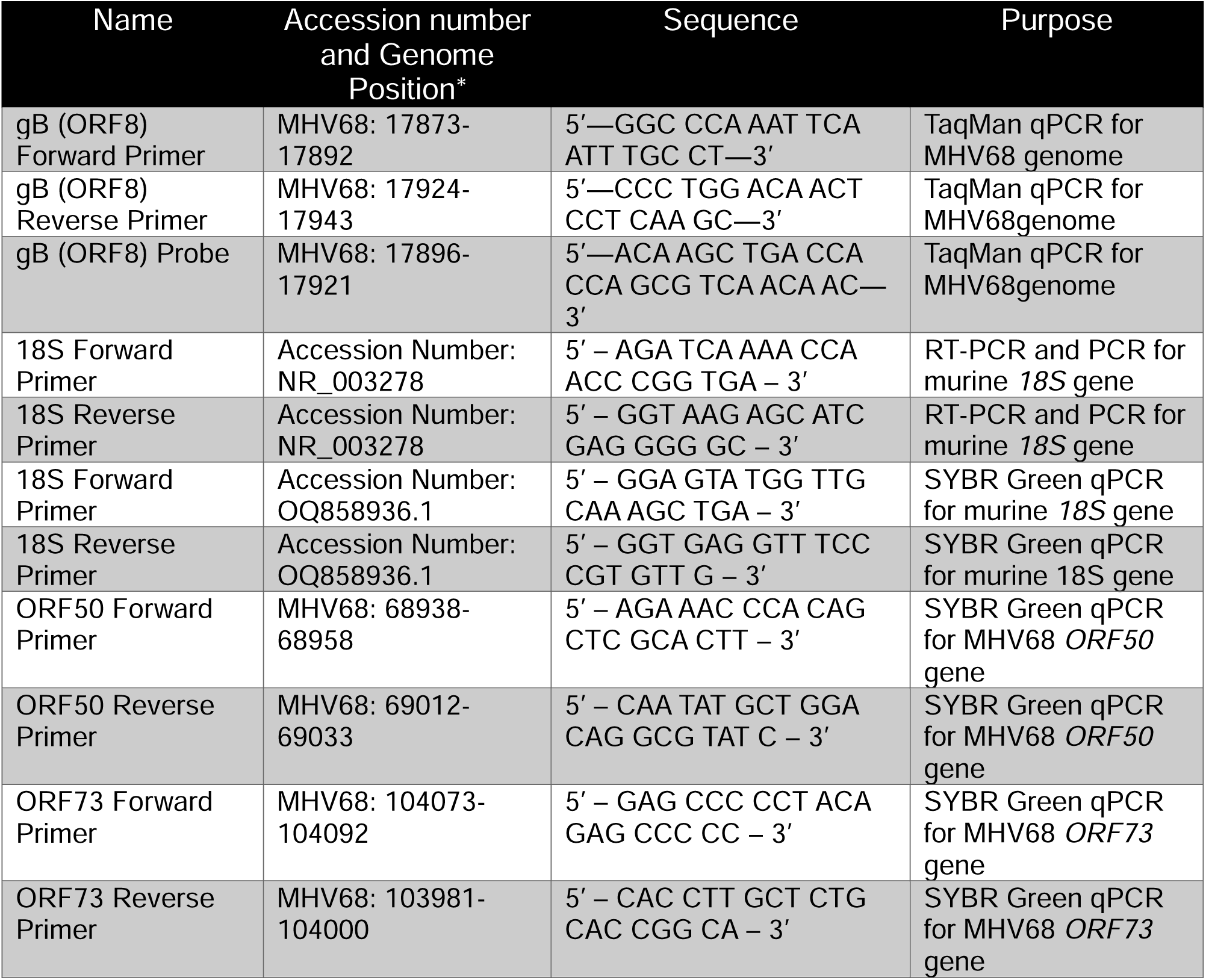
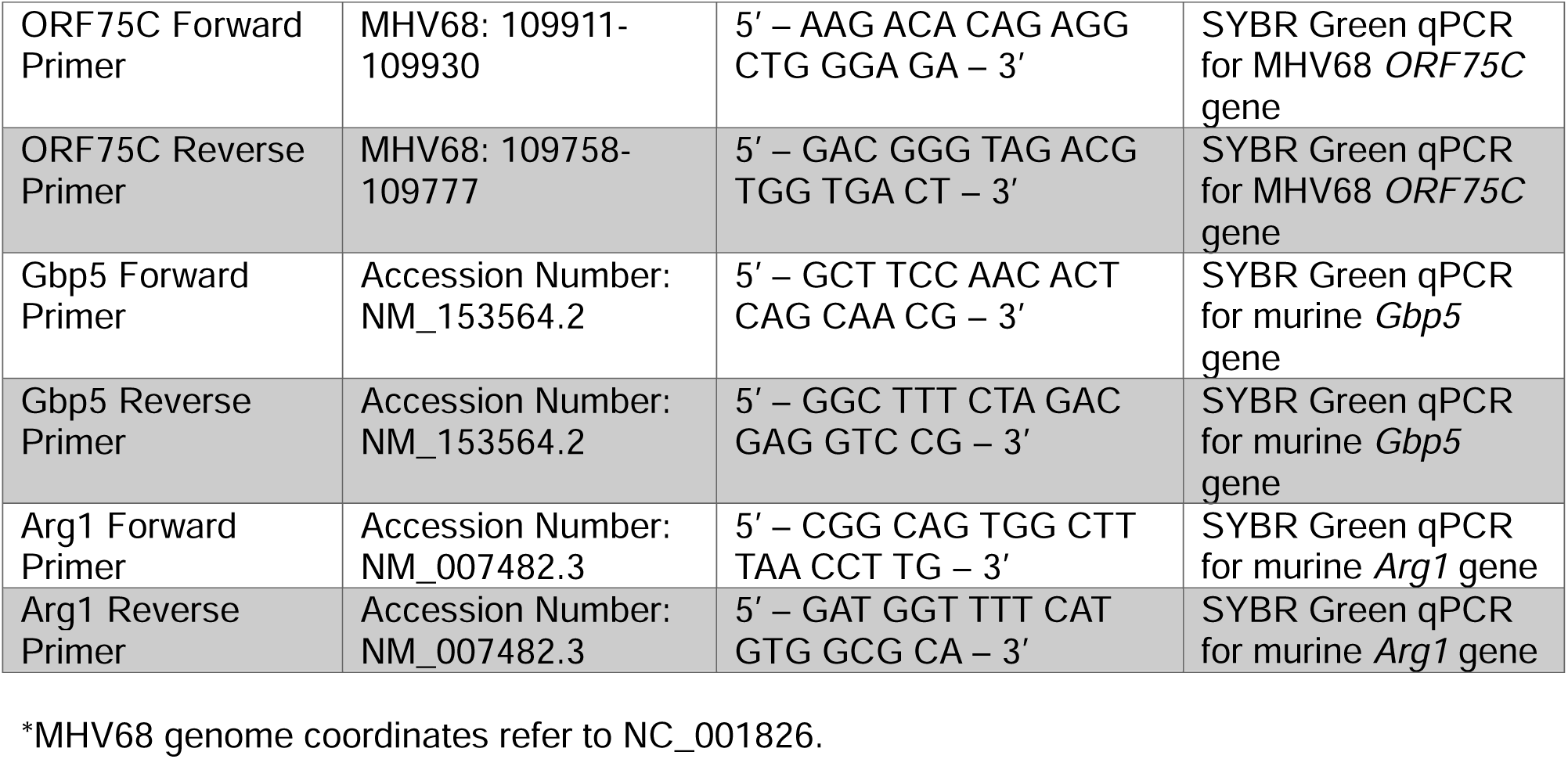
Oligonucleotides for RT-PCR, PCR, and qPCR analysis.

For ex vivo infection of primary peritoneal cells, eight-week old female C57BL/6J mice were euthanized, with peritoneal cells collected via peritoneal lavage, with cells cultured in the presence or absence of IFNγ [100 U/mL]. 16 hours post-plating, cells were infected with indicated virus (MOI=1), with fresh cDMEM ± IFNγ replaced 1 hpi, and samples harvested at 48 hpi.

### Treatment Regimens

The following cytokines and inhibitors and their respective concentrations were used in these studies with commercial sources identified in **Key Resources Table**: interferon gamma, IFNγ [100 U/mL]; interleukin-4, IL-4 [10 ng/mL]; ruxolitinib, RUX [2 µM]; interferon beta, IFNβ [100 U/mL]; SP140 inhibitor, GSK761 [40 nM]; topoisomerase I inhibitor, topotecan [100 nM]; topoisomerase II inhibitor, etoposide [25 µM]. Cells were either treated 16 hours prior to infection (-16 hpi) or 1-hour post-infection (1 hpi); for pre-treatment samples, cultures were replenished with fresh cDMEM ± relevant treatment at 1 hpi. For RUX combination treatments with either IFNγ or IL-4, cells were exposed to RUX for 30 minutes prior to cytokine addition. Treatment regimen concentrations were based on published literature (e.g. IFNγ^18,19,59–61^)

### Virus Attachment and Entry Assays

J774 cells were plated (5x10^5^ cells/well in 6-well plates) with or without IFNγ (16 hr pre-treatment), infected with WT MHV68 virus, incubated at 4°C for 1 hour prior to removal of virus inoculum and washing of cells with cold PBS. For samples collected for virus attachment analysis, PBS wash was removed and replaced with cold cDMEM, with cells scraped, collected, and stored at -80°C. For samples collected for total virus entry, PBS wash was removed and replaced with fresh cDMEM ± IFNγ, incubated for 2 hours at 37°C, after which media was removed, cells were rinsed with warm 0.25% Trypsin-EDTA, replaced with fresh cDMEM, and cells were scraped, collected, and stored at - 80°C. For samples collected for viral nuclear entry, PBS wash was removed and replaced with fresh cDMEM ± IFNγ, incubated for 4 hours at 37°C, after which cells were once again rinsed with cold PBS, replaced with fresh cDMEM, and cells were scraped, collected, and processed for nuclear DNA isolation (detailed in **Supplemental Methods**).

### Flow Cytometry

3T12, J774, or primary peritoneal cells were harvested at indicated time points followed by staining for flow cytometric analysis. Cell suspensions were stained with indicated LIVE/DEAD viability stains at 1:1500 dilution in BSS wash ([111 mM] dextrose, [2 mM] KH_2_PO_4_, [10 mM] NA_2_HPO_4_, [25.8 mM] CaCl_2_·2H_2_O, [2.7 mM] KCl, [137 mM] NaCl, [19.7 mM] MgCl·6H_2_O, [16.6 mM] MgSO_4_), analyzed for various indicators of virus infection, including GFP, vRCA and γH2AX expression, or CCF2-AM or CCF4-AM β-lactamase substrate cleavage. All staining was done in the presence of Fc receptor— blocking antibody (clone 2.4G2), and fixed in 1% paraformaldehyde prior to analysis. For a subset of experiments, to simultaneously quantify total and lytic infection within the same cells, parallel cultures of J774 cells were infected with MHV68.LANA::βlac or MHV68.vCyc::βlac reporter viruses. At time of harvest, cultures with incubated with distinct fluorescently-labelled (BUV395- or AF647-) anti-CD45 antibodies, mixed together, incubated with β-lactamase substrate (CCF2 or CCF4), and analyzed for β-lactamase activity indicative of either LANA.βlac or vCyc.βlac expression. Samples were debarcoded based on expression of either BUV395- or AF647-labelling after sample collection, with the frequency of LANA.βlac or vCyc.βlac expressing cells defined based on the specific fluorescent antibody used to label each input population. To rule out the impact of specific antibody combinations skewing results, experiments reversed fluorescent labels, with no difference observed in the frequency of LANA.βlac or vCyc.βlac expressing cells (not shown). Flow cytometric analysis was performed on an Agilent Novocyte Penteon flow cytometer. All experiments included unstained, single-stain, and full-minus-one controls to define background fluorescence, fluorescent signal spread, and compensation, with WT MHV68 infection used to define background activity for all measurements of β-lactamase cleavage. Gating strategies are illustrated in **Fig S2**, with full experimental details in **Supplemental Methods**.

### Single Cell RNA-sequencing

To characterize MHV68 transcription during primary, acute peritoneal macrophage infection in vivo, female C57BL/6 mice were infected with 1x10^6^ PFU by intraperitoneal injection. Peritoneal cells were harvested and pooled from four infected animals at 16 hpi, stained with live/dead Zombie Near-IR, F4/80-AlexaFluor 647 (clone BM8, 1:200 dilution) in the presence of Fc receptor blockade (clone 2.4G2), and CCF2-AM. Cells were sorted on a Beckman Coulter XDP MoFlo Astrios, purifying live, single, F4/80+ cells that either had CCF2 cleavage (indicating virus infection, defined by the expression of the LANA::βlac fusion protein, i.e. LANA.βlac+ or LANA positive) or lacking CCF2 cleavage (indicating cells that were not infected or failed to express the fusion protein, i.e. LANA.βlac- or LANA negative). Single-cell RNA-seq was prepared using the Chromium 3’ V3 library kit (10X Genomics). Cell processing and sequencing was performed by the University of Colorado Cancer Center Genomics Shared Resource (RRID: SCR_021984), with data processed using Cell Ranger (v6.0.1)^62^ and Seurat (v4.0.4)^56^ (fully detailed in **Supplemental Methods**). Data from LANA.βlac+ cells have been previously reported^12^, with analysis solely focused on viral reads and cell clusters, without comparison to the paired LANA.βlac-sample.

### Statistical Analysis and Software

Data analysis, graphing and statistical analyses were performed using GraphPad Prism (version 11.0.2; GraphPad Software, San Diego, California USA, www.graphpad.com). Flow cytometry data were analyzed using FlowJo (version 10.10.1 Ashland, OR; Becton, Dickinson and Company; 2026). scRNA-seq data processing and analysis were done as described above. Statistical significance was tested by unpaired, nonparametric, Mann-Whitney t test or one-way ANOVA, with statistical test and significance defined in each figure legend. Raw and processed data are available through NCBI GEO (GSE345343).

### Quantitative PCR for MHV68 viral DNA load

MHV68 virus load was detected in DNA samples via primers amplifying *Orf73* and normalized to murine *Fos* locus (**Table 2**), using qPCR with Fast SYBR Green Master Mix (Applied Biosystems). Virus copies were determined with a calibration standard consisting of a synthetic double-stranded DNA template of known sequence and molarity (**Table 2**). Serial PCRs of 2-fold dilutions of standard (std) oligos were run, with oligo copy number ranging from 500,000 copies to 488 copies per PCR. All qPCR was performed on a QuantStudio 7 Pro qPCR system (Applied Biosystems).

**Table 2:**
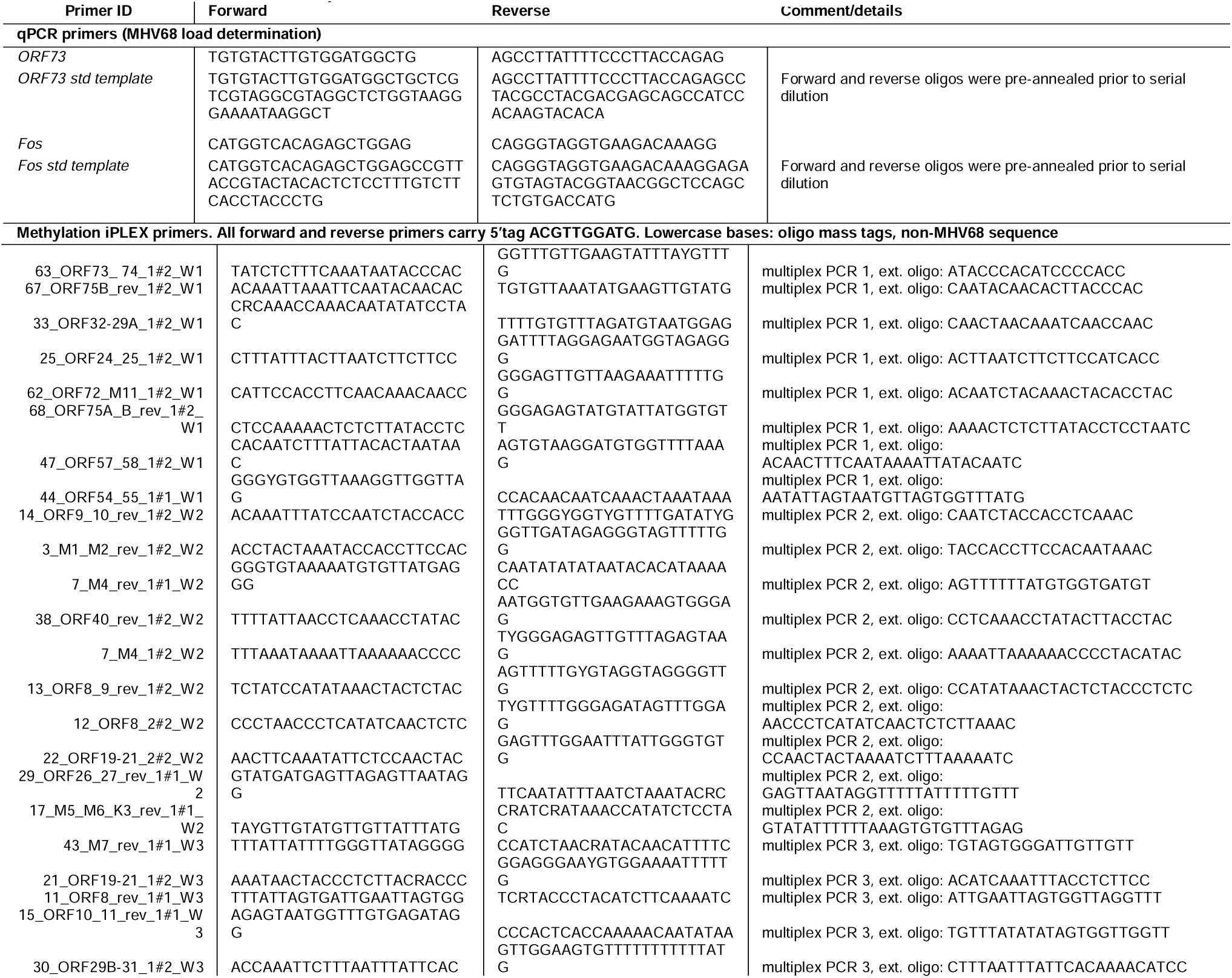
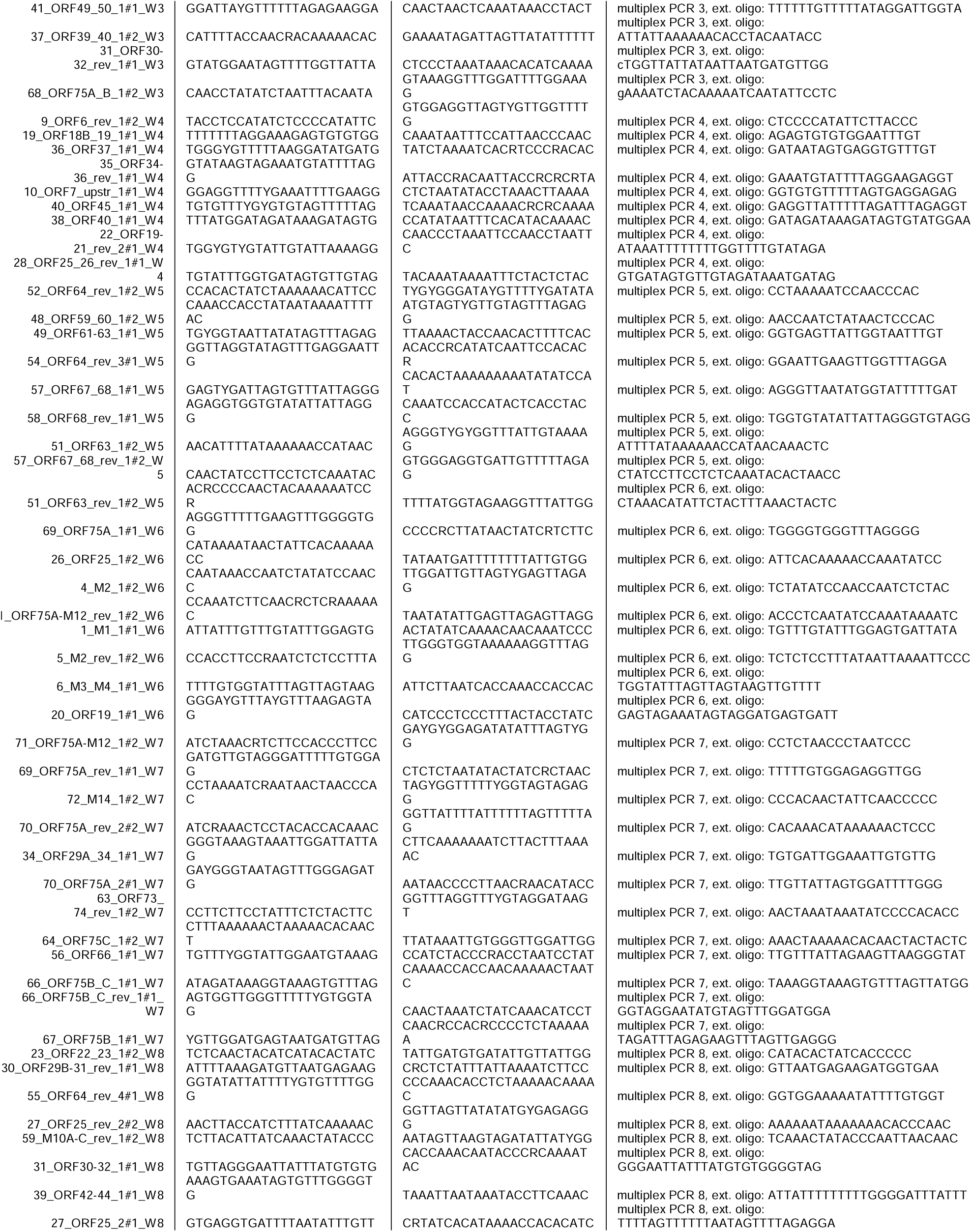
Oligonucleotide primers for iPLEX methylation assay.

### MHV68 DNA methylation assay

MHV68 DNA methylation was measured from 2.0 µg of DNA that was bisulfite converted using EZ DNA methylation kit (Zymo Research) and MHV68 regions were PCR-amplified with primers specific for bisulfite DNA. MHV68 DNA methylation analysis was carried out using the methylation-iPLEX assay as described ^63,64^ (fully detailed in **Supplemental Methods**).

### Ethics Statement

All animal studies were performed in accordance with the recommendations in the Guide for the Care and Use of Laboratory Animals of the National Institutes of Health. Studies were conducted in accordance with the University of Colorado Denver Anschutz Institutional Animal Care and Use Committee (IACUC) under the Animal Welfare Assurance of Compliance policy (assurance no. D16-00171). All procedures were performed under isoflurane anesthesia, and all efforts were made to minimize suffering.

## SUPPLEMENTAL FIGURE LEGENDS

**Supplemental Figure 1 (Fig S1).**
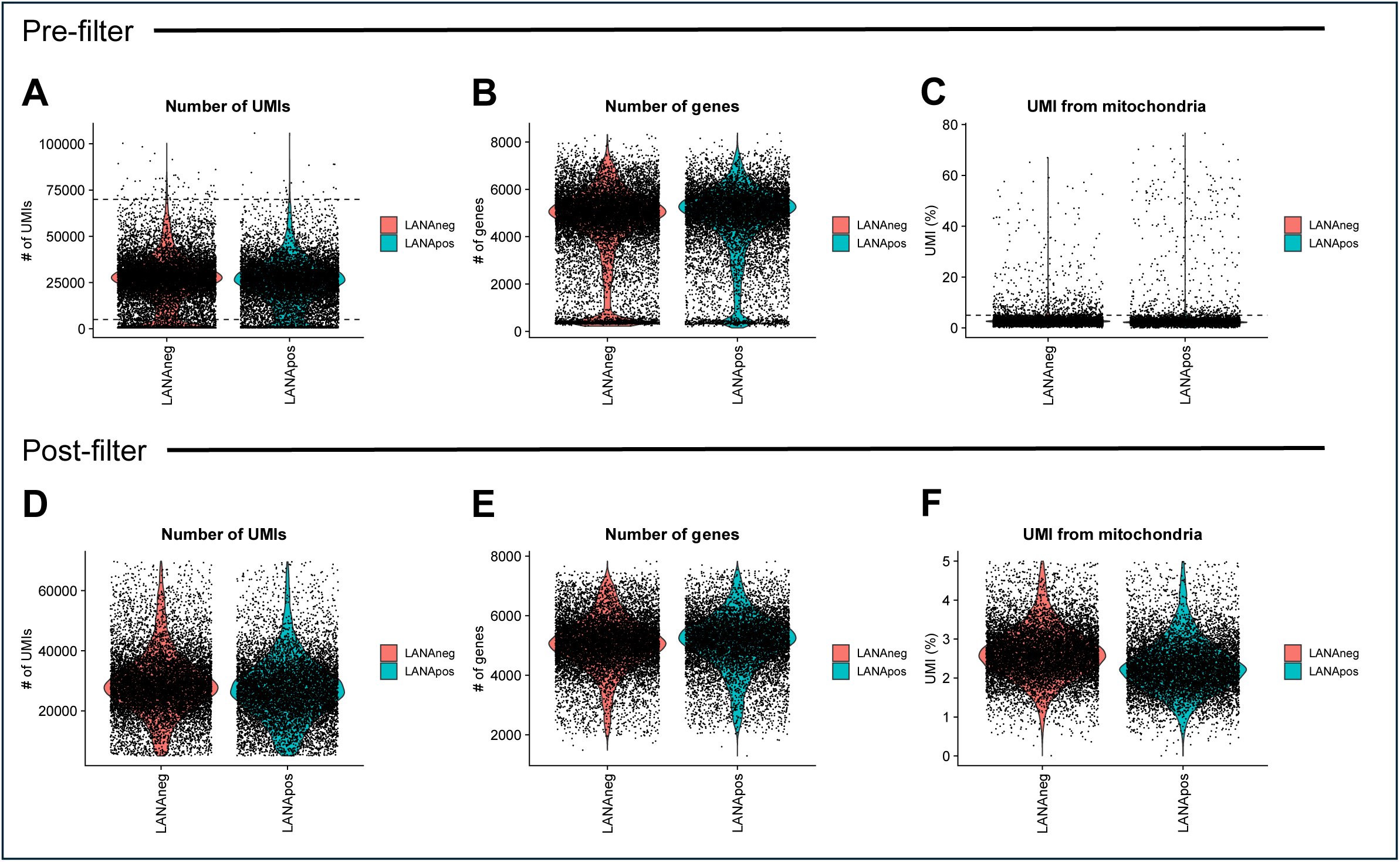
Pre- and post-filtering quality control metrics for scRNA-seq analysis. Data correspond to Figure 1. Quality control metrics of scRNA-seq of primary peritoneal macrophages sort-purified from MHV68-infected C57BL/6J mice at 16 hpi. Violin plots, with individual per cell values depicted by individual circles, show **(A-C)** data before or **(D-F)** after implementing filters. Horizontal dashed lines in panels A-C represent values for cutoffs applied to generate panels D-F. Data depict **(A,D)** number of UMIs per cell, **(B,E)** number of genes per cell, and **(C,F)** percent of UMIs derived from mitochondria. Post-filter datasets contain 11,396 LANAβlac-cells and 9,537 LANAβlac+ cells.

**Supplemental Figure 2 (Fig S2):**
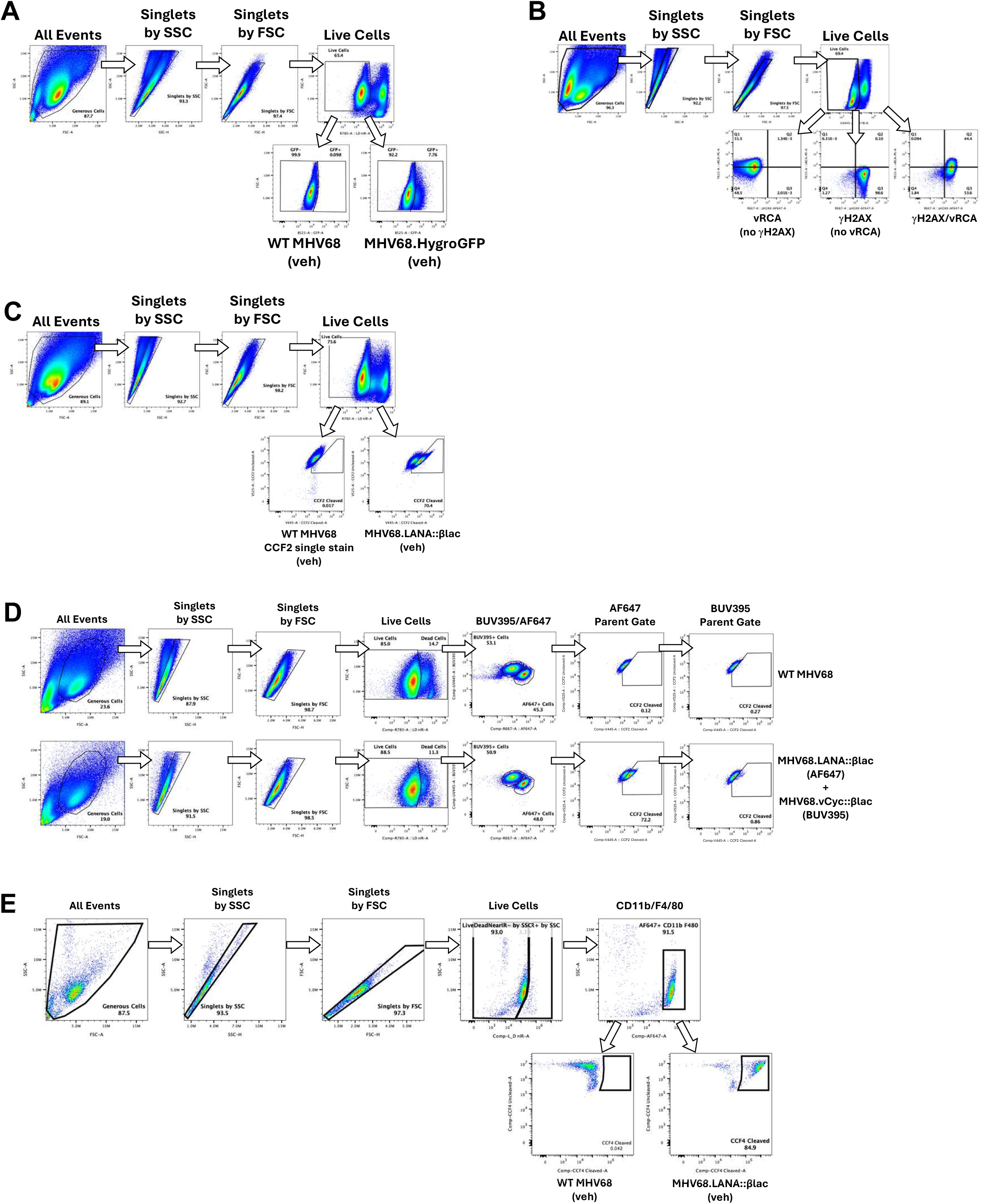
Gating strategies for flow cytometric analysis of MHV68 infection of macrophages. Data correspond to Figures 2**, 3, 5, 6 and 7 and Figure S3**. Analysis of viral gene expression in MHV68-infected **(A-C)** J774 cells or **(D-E)** primary peritoneal cells by flow cytometry. **(A-E)** Gating strategy to analyze MHV68-infected J774 cells **(A-C)** or primary peritoneal cells **(D, E)**. In all cases, infected cells were analyzed by flow cytometry, using sequential focused analysis (i.e. “gating”) to define single cells (“Singlets by FSC” and “Singlets by SSC”) that are viable (“Live Cells”) by exclusion of a dye that positively identifies dead cells. **(A)** In support of Figures 2B**-2G, 3A** and **Figure S3**, single, viable J774 cells were analyzed for GFP fluorescence, with WT MHV68-infected cells (which lack GFP) used to define background fluorescence relative to MHV68.HygroGFP-infected cells. **(B)** In support of Figure 5A-B, gating strategy to analyze vRCA and γH2AX expression in WT or C-RTA MHV68-infected J774 cells. Viable, single cells were analyzed for antibody binding for lytic protein markers, with positive populations further informed by control samples (including single stain and full-minus-one controls). Representative data shown from MHV68.C-RTA infected samples. **(C)** In support of Figures 6A, single, viable J774 cells (J774s) infected with an MHV68 β-lactamase reporter virus were gated into substrate uncleaved or cleaved populations based on distinct fluorescence emission, with WT MHV68-infected cells (which lack beta-lactamase) used to define background fluorescence, above which cells were considered to have cleaved substrate fluorescence. **(D)** In support of Figure 7A-E, where single, viable cells infected with an MHV68 β-lactamase reporter virus were separated into distinct populations based on fluorescence from an anti-CD45 antibody binding (BUV395 or AF647; dual-reporter system). BUV395+ or AF647+ populations were subsequently analyzed for the frequency of substrate cleaved events as an indicator of β-lactamase events (either LANA.βlac+ to define total infection, including latent and lytic infection gene expression or vCyc.βlac+ events to define cells with lytic gene expression). **(E)** In support of Figure 7F-H, single, viable, ex vivo primary peritoneal cells infected with either WT MHV68 (bottom row, left) or MHV68 β-lactamase reporter virus (bottom row, right) were identified as macrophages based on expression of CD11b and/or F4/80, after which cells were analyzed for beta-lactamase substrate cleavage. WT MHV68-infected cells (which lack beta-lactamase) were used to define background fluorescence, above which cells were considered to have cleaved substrate fluorescence indicative of beta-lactamase expression.

**Supplemental Figure 3 (Fig S3):**
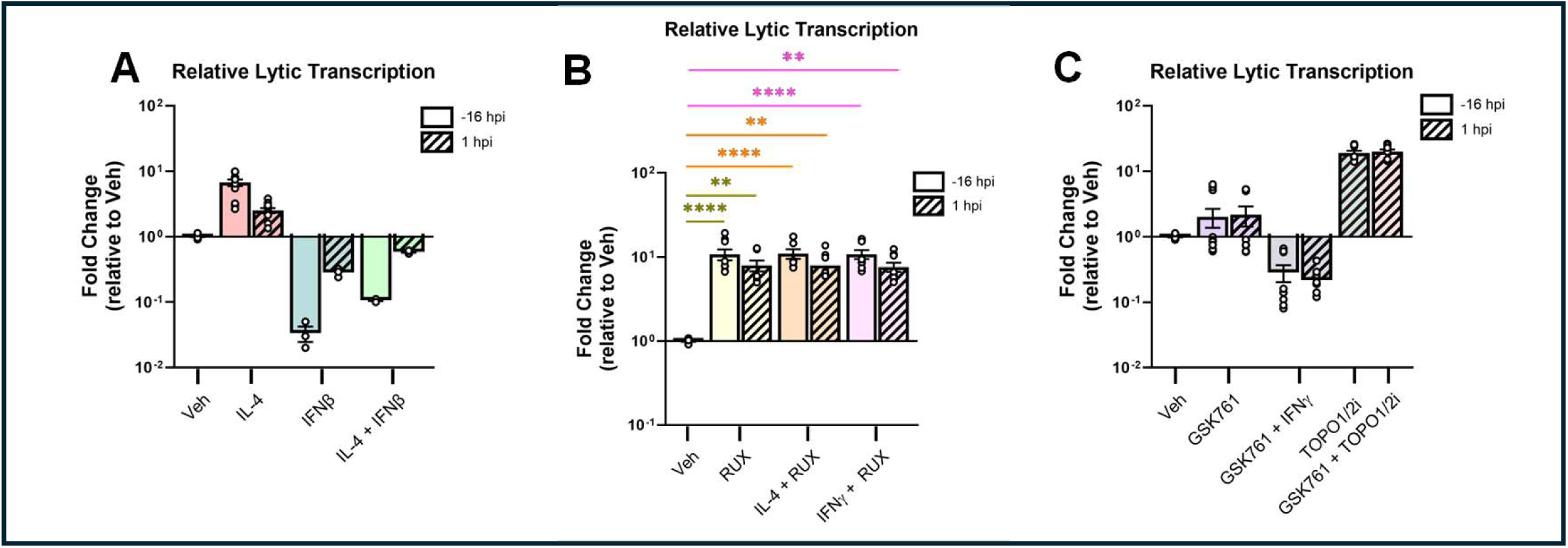
Effect of cytokines, pharmacological inhibitors, and combinatorial treatments on MHV68 lytic transcription in J774 macrophages. Data correspond to Figure 3. Analysis of viral gene expression in MHV68.HygroGFP-infected J774 myeloid cells (MOI=1 PFU/cell, harvested at 48 hpi) treated with the indicated conditions before (-16 hpi, open bars) or after (1 hpi, diagonal bars) infection. **(A-C)** Quantitation of the impact of (A) IL-4, IFNβ or the combination, (B) ruxolitinib (RUX) alone or in combination with IL-4 or IFNγ, or (C) GSK761 (Sp140 inhibitor), TOPO1/2i (topoisomerase inhibitor cocktail, Etoposide + Topotecan), or the indicated combinations. Plots depict the fold change in the frequency of GFP+ cells among single, viable cells (gating strategy in **Fig S2A**) relative to vehicle controls (defined as 1). Data from **(A)** 1 (IFNβ) or 3 independent experiments (Veh, IL-4), **(B)** 3 independent experiments, or **(C)** 2 independent experiments; in all cases, experiments contained 3 technical replicates per sample, showing mean ± SEM, with individual symbols depicting all replicate samples. Data for **(A)** IL-4 and **(B)** RUX treated samples are the same data shown in Figure 3A, as these samples were matches controls for **(A)** IL-4+IFNβ and **(B)** IL-4+RUX and IFNγ+RUX. Statistical analysis (**B)** was done using one-way ANOVA (non-parametric, Kruskal-Wallis with Dunn’s multiple comparisons), comparing vehicle to each treatment type in individual statistical tests. Statistical significance is indicated as \*\**p* <0.01, \*\*\*\*\**p* <0.0001. Veh, Vehicle; IL-4, Interleukin 4; IFNβ, interferon beta; RUX, ruxolitinib; GSK761, an Sp140 inhibitor; TOPO1/2i, topoisomerase inhibitor cocktail, Etoposide + Topotecan.

**Supplemental Figure 4:**
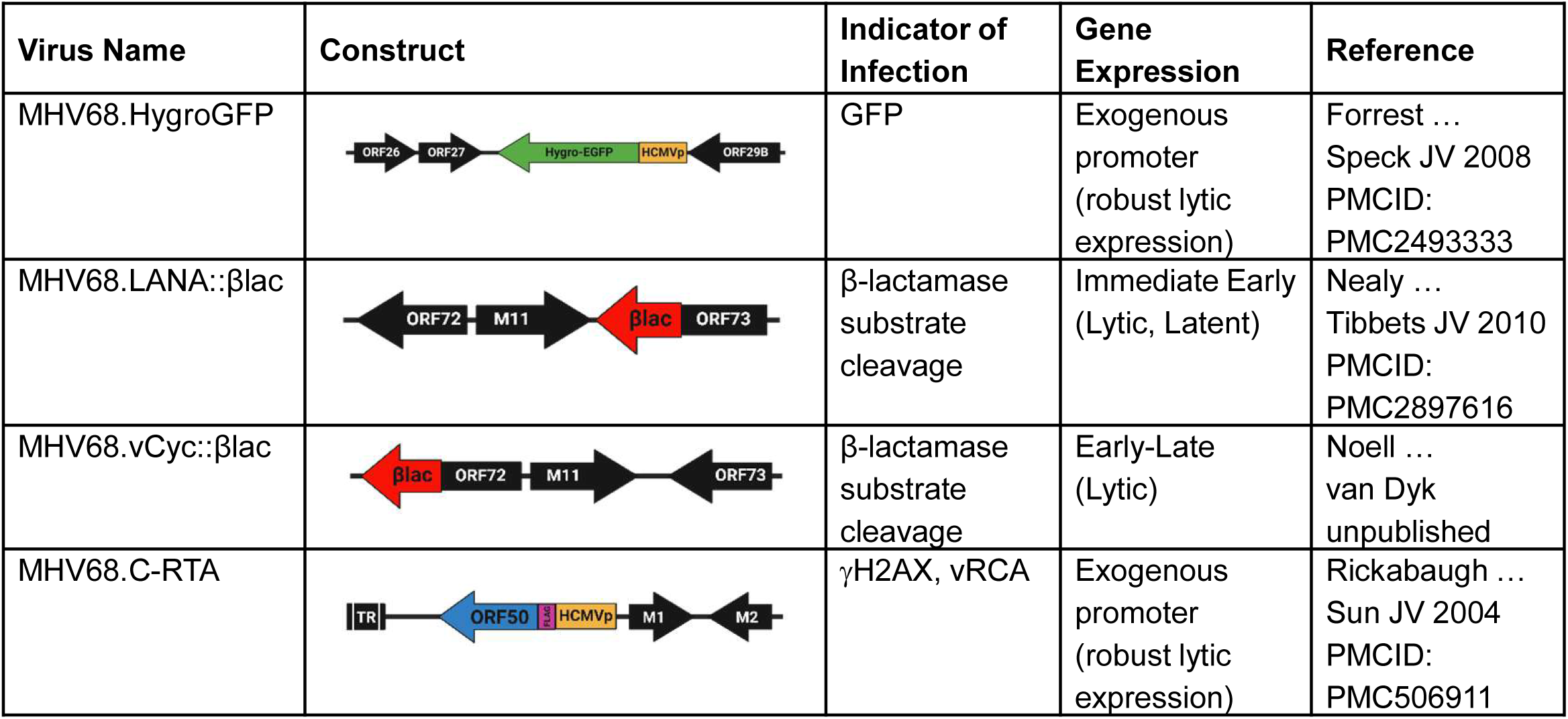
MHV68 recombinant reporter virus constructs. Recombinant MHV68 viruses used in the described work. **MHV68.HygroGFP** has an exogenous hygromycin eGFP gene transcribed by the human cytomegalovirus immediate-early promoter (HCMVp) which robustly expresses GFP during lytic infection. **MHV68.LANA::**β**lac** encodes an *Orf73*::β-lactamase gene fusion, resulting in expression of a protein fusion between LANA and β-lactamase. This gene is transcribed with immediate early kinetics and expressed during both latent and lytic infection. **MHV68.vCyc::**β**lac** encodes an *Orf72*::β-lactamase gene fusion, resulting in expression of a protein fusion between the MHV68 viral cyclin (vCyc) and β-lactamase. This gene is transcribed with early-late kinetics, expressed during lytic infection. **MHV68.C-RTA** encodes a flag-epitope tagged *Orf50*, which encodes the RTA protein, inserted at the left end of the MHV68 genome. RTA expression is driven by the human cytomegalovirus immediate-early promoter (HCMVp), resulting in constitutive expression of this lytic cycle viral transactivator. This virus is latency deficient and is characterized by enhanced lytic replication. Lytic replication for C-RTA (and WT MHV68 infection) is quantified by measuring expression of two protein modifications associated with lytic replication, γH2AX phosphorylation and vRCA expression. Constructs are based on published references for each recombinant as identified.

