## Supplemental Methods for "Cytokine signaling regulates multiple stages of gammaherpesvirus infection in myeloid cells"

**Plaque Assay**

Plaque assay quantification of viral titer was performed using mouse 3T12 fibroblast cells (ATCC CCL-164). Cells were plated in 12-well plates at 8.5x10^4^ cells per well one day prior to infection. Viral samples were diluted 10-fold in 5% cDMEM. An internal standard was included for each assay to ensure reproducible sensitivity for each plaque assay. Cells were incubated with virus for 1 hour at 37^o^C at 5% CO_2_. Plates were rocked every 15 min. Cells were then covered by an overlay composed of a 1:1 mix of 10% cDMEM and carboxymethyl cellulose (CMC; Sigma, Cat. No. C-4888) supplemented with Gibco Amphotericin B (Thermo Fisher Scientific, Cat. No. 15290018). Cells were incubated for 8 days before staining with 0.5% methylene blue and plaques were counted.

**Nuclear DNA Isolation**

Cells collected were pelleted by centrifugation for 5 minutes at 500xg at 4^o^C, resuspended in 250 µL ice-cold cell membrane lysis buffer ([320 mM] sucrose, [10 mM] HEPES, [8 mM] MgCl_2_, [1X] Protease/Phosphatase Inhibitor Cocktail, and [0.01%] v/v Triton X-100), and incubated on ice for 15 minutes^1^. Cells were pelleted again by centrifugation for 5 minutes at 500xg at 4^o^C, and the supernatant containing the cytosolic fraction was removed and discarded. Cells were then washed twice with 250 µL nuclear lysis buffer ([320 mM] sucrose, [10 mM] HEPES, [8 mM] MgCl_2_, and [1X] Protease/Phosphatase Inhibitor Cocktail). Samples were pelleted by centrifugation for 5 minutes at 500xg at 4^o^C, and nuclear pellets were resuspended in 400 µL phenol:chloroform:isoamyl alcohol, followed by vortexing. 400 µL of 1:9 TES buffer ([10%] SDS, [1X] TE buffer) was added to each sample followed by vortexing and a 30-minute incubation at 56^o^C. Samples were pelleted by centrifugation for 10 minutes at max speed (22,000x g) at room temperature. Aqueous layers were collected and DNA was followed by the addition of 400 µL chloroform and mixed by gentle inversion. Samples were pelleted by centrifugation for 10 mins at max speed at room temperature, aqueous layers were once again collected and DNA was precipitated by the addition of 40 µL of 3M sodium acetate, and 880 µL of cold 100% EtOH followed by gentle mixing and an overnight incubation at -20^o^C. After incubation, samples were pelleted by centrifugation at max speed for 20 minutes at room temperature. Supernatants were discarded and pellets were washed with 200 µL of 70% EtOH. Samples were spun down at max speed for 5 minutes at room temperature, supernatants discarded, and pellets air dried at room temperature. Once dry, DNA pellets were resuspended in 30 µL of molecular grade water.

**Quantitative PCR for viral DNA**

Infected cells were harvested by scraping at indicated harvest time (hpi), subjected to three freeze/thaw cycles, with DNA isolated using an overnight incubation in a 1:1:8 solution of 10% SDS : Proteinase K [10 mg/mL] : 1X TE buffer, respectively, at 56^o^C for heat inactivation. DNA was normalized to a concentration of 20 ng/µL in molecular grade water. qPCR analysis was done with 100 ng of DNA using a LightCycler 480 Probe Master-Mix kit (Roche, Cat. No. 04707494001) and a primer and probe set specific to MHV68 gB to quantify the number of viral genome copies. Primers and probes listed in **Table 1**. A glycoprotein B (gB) standard curve was generated using a gB plasmid dilution series ranging from 10^2^ to 10^10^ copies diluted in background DNA, with a limit of detection (LOD) of 100 copies.

**RT-qPCR**

RNA was isolated at indicated times by 10-minute incubation in TRIzol reagent, followed by TURBO DNase treatment according to manufacturer’s protocols. RNA amplification and removal of DNA was confirmed by PCR amplification of the control host gene, 18S, in the presence or absence of reverse transcriptase. RT-PCR was performed using the OneStep RT-PCR kit (Qiagen) with the following conditions: (i) 50^o^C for 30 min, (ii) 95^o^C for 15 min, (iii) 40 cycles of 94^o^C for 30 sec, 52^o^C for 30 sec, and 72^o^C for 30 sec, (iv) 72^o^C for 10 min, and (v) hold at 4^o^C. DNA-free RNA samples were converted to cDNA using random primers (250 ng/µL) and Superscript II reverse transcriptase following the manufacturer’s protocol. 100 nanograms of cDNA was used for qPCR analysis of the identified genes using the iQ SYBR green supermix with the following conditions: (i) 95^o^C for 3 min, (ii) 40 cycles of 95^o^C for 15 sec, 60^o^C for 1 min, and (iii) 95^o^C for 15 sec, 60^o^C for 1 min, and 95^o^C for 15 sec. Amplification of viral genes was normalized to murine 18S expression to calculate the relative difference of the target gene expression using the Pfaffl method, as previously^2,3^ described: Target primer efficiency^TargetΔ*Ct*/18S primer efficiency^18SΔ*Ct*. A single product for each target was confirmed by melt curve analysis. PCR primers are listed in **Table 1**.

**Flow Cytometry**

3T12, J774, or primary peritoneal cells were harvested at indicated time points followed by staining for flow cytometric analysis. Cell suspensions were stained with indicated LIVE/DEAD viability stains at 1:1500 dilution in BSS wash ([111 mM] dextrose, [2 mM] KH_2_PO_4_, [10 mM] NA_2_HPO_4_, [25.8 mM] CaCl_2_·2H_2_O, [2.7 mM] KCl, [137 mM] NaCl, [19.7 mM] MgCl·6H_2_O, [16.6 mM] MgSO_4_). Cells were then either stained with CCF2-AM or CCF4-AM β-lactamase substrate (due to extensive product backorder of CCF2-AM reagent, CCF4-AM was used) or antibody stained for protein detection. Cells were incubated in 1:10 (in vitro cell lines) dilution or neat (primary cells) β-lactamase substrate (CCF2 or CCF4) following the manufacturer’s protocol. Cells were then stained with fluorescently conjugated antibodies to ORF4 (a rabbit antibody against the MHV68 ORF4 protein (vRCA), a gift of the Virgin Lab, Washington University St. Louis)^2,4^ (1:400) labeled with a Zenon R-phycoerythrin anti-rabbit IgG reagent, γH2AX (1:800), CD45 (1:400), F4/80 (1:400), CD11b (1:400). All staining was done in the presence of Fc receptor—blocking antibody (clone 2.4G2), and fixed in 1% paraformaldehyde prior to analysis. Flow cytometric analysis was performed on an Agilent Novocyte Penteon flow cytometer. All flow cytometry experiments included unstained, single-stain, and full-minus-one controls to define background fluorescence, fluorescent signal spread, and compensation.

**Dual Reporter System (LANA::βlac and vCyc::βlac co-detection)**

J774 cells were plated in 6-well plates at a density of 5x10^5^ cells/mL in cDMEM ± vehicle or ± IL-4 [10 ng/mL] and plates were incubated overnight at 37^o^C with 5% CO_2_. 16 hours post-plating, cells were infected with virus inoculum at an MOI=1 PFU/cell and incubated for 1 hour at 37^o^C and 5% CO_2_. After incubation, virus inoculum was removed and replaced with fresh cDMEM containing vehicle treatment or IL-4 and cells were incubated for 24 or 48 hours post infection at 37^o^C and 5% CO_2_. 24- and 48-hour samples were staggered to allow cells for both time points to be harvested and processed at the same time. At time of processing, cells were scraped and collected with supernatants into tubes and kept on ice. Samples were pelleted by centrifugation for 5 minutes at 500xg at 4^o^C, supernatant was removed and discarded, then cell pellets were resuspended in 1 mL ice-cold BSS buffer. All samples were kept on ice throughout protocol with the exception of room temperature antibody incubations.

Samples were stained with CD45 antibodies conjugated to either AlexaFluor 647 or BUV395 depending on the virus assignment to fluorophore (e.g. MHV68.LANA::βlac virus assigned to AF647 and MHV68.vCyc::βlac virus assigned to BUV395, and vice versa in a separate experiment to eliminate virus-fluorophore artifacts). Samples were pelleted by centrifugation for 5 minutes at 500xg at 4^o^C, followed by pellet resuspension in 100 µL of appropriate sample-specific CD45 antibody. Samples were incubated for 30 minutes in the dark at room temperature. Following incubation, antibody stains were diluted with 900 µL BSS buffer. Samples that were to be combined to detect populations of total infected and lytic infected cells were pelleted by centrifugation for 5 minutes at 500xg at 4^o^C. Supernatants were removed and discarded, and sample combination pair pellets were resuspended together in a total of 1 mL BSS buffer.

Following CD45 staining, individual and combined samples were pelleted by centrifugation for 5 mins at 500xg at 4^o^C, followed by resuspension of pellet in 100 µL of 1:1000 Live/Dead NearIR viability dye in BSS buffer. Samples were incubated for 30 minutes in the dark at room temperature. Following incubation, LIVE/DEAD stain was diluted by adding 900 µL to each sample.

Following live/dead staining, individual and combined samples were pelleted by centrifugation for 5 minutes at 500xg at 4^o^C. BSS supernatants were discarded and pellets were resuspended in 200 µL non-additive DMEM (naDMEM). 40 µL of beta-lactamase loading solution containing CCF2 substrate was added to each sample. Samples were incubated for 30 minutes in the dark at room temperature with vortexing of tubes every 10 minutes. CCF2 stain was then diluted by adding 300 µL naDMEM to each sample. To wash cells of any remaining substrate solution, samples were pelleted by centrifugation for 5 minutes at 500xg at 4^o^C, supernatant was removed and discarded, and pellets were resuspended in 200 µL naDMEM. All samples were pelleted by centrifugation for 5 minutes at 500xg at 4^o^C. Supernatants were removed and discarded, and cell pellets were fixed by resuspension in 200 µL 1% PFA. Samples were then transferred to a 96-well plate for flow cytometric analysis.

**Single Cell RNA-sequencing**

To characterize MHV68 transcription during primary, acute peritoneal macrophage infection in vivo, female C57BL/6 mice were infected with 1x10^6^ PFU by intraperitoneal injection. Peritoneal cells were harvested and pooled from four infected animals at 16 hpi, stained with live/dead Zombie Near-IR, F4/80-AlexaFluor 647 (clone BM8, 1:200 dilution) in the presence of Fc receptor blockade (clone 2.4G2), and CCF2-AM. Cells were sorted on a Beckman Coulter XDP MoFlo Astrios, purifying live, single, F4/80+ cells that either had CCF2 cleavage (indicating virus infection, defined by the expression of the LANA::βlac fusion protein, i.e. LANA.βlac+ or LANA positive) or lacking CCF2 cleavage (indicating cells that were not infected or failed to express the fusion protein, i.e. LANA.βlac- or LANA negative). Data from LANA.βlac+ cells have been previously reported^5^, with analysis solely focused on viral reads and cell clusters, without comparison to the paired LANA.βlac- sample. Single-cell RNA-seq was prepared using the Chromium 3’ V3 library kit (10X Genomics). Cell processing and sequencing was performed by the University of Colorado Cancer Center Genomics Shared Resource (RRID: SCR_021984). Resulting data were processed as follows: Cell Ranger (v6.0.1)^6^ was used to process the fastq files to cell and gene count tables using unique molecule identifiers (UMIs) with the include-introns parameter. Because of the difficulties counting viral reads, this was performed in a two-pass manner. In the first pass, reads were aligned to a chimeric genome of mouse mm10 (GENCODE M23 gene annotations) and MH636806.1 (no gene annotations, only positive and negative strand alignment). The viral counts were stored in metadata and removed from the counts matrix. Reads mapping to MH636806.1 in step were aligned to the same chimeric genome but in this case, the viral transcriptome as well as specific intergenic regions were annotated. The intergenic regions were determined as the strand-specific sequences between genes with 1 base pair of padding on both ends. To minimize reads overlapping with multiple viral genes, only the first 70bp of each read were aligned and counted. The resulting counts matrix of viral alignment was appended to the host gene counts from step 1.

The Seurat (v4.0.4)^7^ pipeline was used for downstream quality control and analysis. Cell Ranger-filtered data was read into Seurat. Host genes were removed if identified in fewer than 10 cells, while viral gene and intergenic regions were removed if found in no cells. Cells were removed if they expressed <50 genes or viral regions, <5000 UMIs, >5% total UMIs from mitochondria, or >70000 UMIs. The filtered data was normalized by dividing gene counts by total counts per cell and multiplied by 10,000 followed by natural-log transformation. The top 2,000 most variable genes were scaled with total UMI and percentage mitochondria regressed out. These were used to calculate principal components (PCs) with the top 20 used to perform Uniform Manifold Approximation and Projection (UMAP) and determining the k-nearest neighbors and clustering.

Clusters were identified by over-representation analysis examining the top enriched markers. Canonical cell type markers were plotting in UMAP space and using dot plots. Dot plots, bar graphs, scatter plots, and violin plots were generated using Seurat and ggplot2 R packages (v3.4.4)^8^.

**MHV68 DNA methylation assay**

MHV68 DNA methylation was measured from 2.0 µg of DNA that was bisulfite converted using EZ DNA methylation kit (Zymo Research) and MHV68 regions were PCR-amplified with primers specific for bisulfite DNA. MHV68 DNA methylation analysis was carried out using the methylation-iPLEX assay as described ^9,10^. In short, iPLEX capture and extension primers were designed from bisulfite-converted DNA sequences using the Typer4.0 software (Agena Bioscience, **Table 2**) and assembled in 8 multiplex PCR reactions to cover n=73 CpG sites across the MHV68 genome. After amplification, dNTPs were removed with Shrimp alkaline phosphatase treatment (Agena Bioscience) for 30 minutes at 37 degrees, followed by heat inactivation at 85 degrees for 5 minutes. Targeted cytosines in CpG dinucleotides then underwent multiplexed single-base extension (iPLEX gold kit, Agena Biosciences) onto the cytosine of interest to measure a quantitative ratio of unmethylated/methylated CpG. Base-extended products were analyzed via Matrix-Assisted Laser Desorption/Ionization-Time of Flight (MALDI-ToF) mass spectrometry (Agena Bioscience). Ratios of unmethylated versus methylated mass peaks were used to calculate the percentage of DNA methylation. Samples were dispensed onto 384-well SpectroCHIP arrays using the RS1000 Nanodispenser and analyzed using the MassARRAY Analyzer4 system (Agena Bioscience). All samples were analyzed in technical duplicates, with average methylation across duplicate readouts in n=73 CpG dinucleotide sites (**Table 2**).
